# Hypoxia-activated prodrug and chemotherapy disrupt resistance-associated metabolism in osteosarcoma

**DOI:** 10.64898/2026.06.16.732534

**Authors:** Sophie M. Pearce, Neil A. Cross, Lucy E. Flint, Malcolm R. Clench, David P. Smith, Daniel M. Allwood, Bradley J. Wallace, Joseph D. Ready, Gregory Hamm, Richard J.A. Goodwin, Laura M. Cole

## Abstract

Chemotherapy resistance remains a critical barrier in treating osteosarcoma. Hypoxia-activated prodrugs (HAPs) target oxygen-deprived tumor regions that evade conventional chemotherapy. Here, we applied integrated spatial multimodal mass spectrometry imaging of metabolites (DESI-MSI, MALDI-MSI), targeted proteomics (IMC), and metallomics (LA-ICP-MSI) to naïve and newly developed doxorubicin-resistant osteosarcoma spheroids treated with a novel HAP tirapazamine analogue, TPZ-A-X, and doxorubicin. Combination treatment selectively downregulated GLUT1 and suppressed pro-survival pAkt in doxorubicin-resistant spheroids whilst inducing comparable DNA damage (γH2AX) across both phenotypes. Metabolomics imaging identified ferroptosis pathway suppression in doxorubicin resistance, which combination treatment reversed, whilst simultaneously depleting glycolytic fuels. Integrative protein-metabolite correlation analysis uncovered functional couplings between glucose transport and CoA-dependent metabolism and spatially revealed anabolic signaling at spheroid peripheries. Combination treatment induced endogenous copper, zinc and magnesium depletion, independent of ATP/ADP collapse reflecting an adaptive survival remodeling of the metalloproteome. HAP/chemotherapy combinations exploit metabolic vulnerabilities via coordinated disruption of ferroptosis suppression, glycolytic dependence, and survival pathways underlying apoptotic resistance. These findings demonstrate a framework for informing mechanistic reasoning and combination strategy design in chemotherapy-resistant tumors.

## INTRODUCTION

High attrition rates in drug discovery continue to present significant challenges to efficiency across the pharmaceutical industry, with a large proportion of candidate drugs failing only after substantial investment into late-stage development. The recently published FDA 2025 roadmap for new approach methodologies (NAMs) emphasizes the need to make animal studies the exception rather than the rule^1,2^, urging a strategic shift toward advanced human-relevant *in vitro* models capable of identifying ineffective candidates earlier in the discovery pipeline, thereby improving the translational relevance and predictive accuracy of early drug discovery. Three-dimensional (3D) multicellular tumor spheroid (MCTS) models^3,4^, used in the work presented here, partially address this need by reproducing key features of the tumor microenvironment absent in traditional 2D monolayer culture, including oxygen and nutrient gradients, endogenous extracellular matrix signaling, and microenvironment-dependent drug resistance mechanisms^5^. Whilst spheroid models remain less complex than more resource-intensive physiologically advanced models, their human-cell-derived origin and capacity to recreate an intrinsic 3D microenvironment position them as strategically valuable intermediaries. The ability to ’fail fast’ using translationally relevant 3D models, which can integrate into routine discovery workflows at strategically advantageous stages, represents an opportunity to improve predictive accuracy of therapeutic response, whilst reducing reliance on animal testing and enhancing overall drug discovery efficiency.

This challenge is particularly critical for osteosarcoma, the most common primary bone malignancy in children and adolescents^6–8^, where five-year survival rates for metastatic and recurrent disease remain below 30%^9,10^. Standard of care chemotherapy relies on high-dose doxorubicin (Dox), cisplatin, and methotrexate combinations^11^, but resistance to these agents is frequently acquired and subsequent prognosis following relapse is poor^12,13^. Our previous work has demonstrated that osteosarcoma 3D cell culture models mimic key features of *in vivo* solid tumors, with important implications for understanding treatment responses^14^. Characterization of ‘aggregoid’ 3D cell culture tumor models revealed heterogeneous metabolic phenotypes with distinct regions of hypoxia and proliferation^15,16^, and comparison against patient tissue architecture identified similarities in metabolic behavior ^17^. Analysis of OS models exposed to acute Dox treatment revealed how the spheroid microenvironment critically shapes the endogenous molecular response, shown by distinct intratumoral remodeling patterns^14^. Building on these findings, the present study extends the investigation to the mechanisms driving chemotherapy resistance, using newly developed Dox-resistant (DXR) osteosarcoma cells, generated through inducing long term acquired resistance, compared against treatment-naïve counterparts. Since the first reported multidrug-resistant human osteosarcoma cell lines were developed^18^, a range of resistant variants have been produced and studied to varying degrees^19^. Using our resistant models, by in depth mapping of molecular signatures using multimodal mass spectrometry imaging (MSI)^20,21^, here specifically, we investigated previously un-characterized responses to hypoxia-activated prodrug (HAP) and chemotherapy combination treatments across both treatment-naïve and drug-resistant models. This work was therefore carried out to provide mechanistic insight into how resistant cells evade therapeutic pressure, to inform rational strategies for amenable targeting of metabolic vulnerabilities and discover biomarkers predictive of therapeutic response in chemotherapy-resistant osteosarcoma.

Hypoxia plays a central role in driving chemotherapy resistance by enabling cells in oxygen-deprived regions to evade cytotoxic agents, reoxygenate, and re-enter the cell cycle to sustain tumor progression. HAPs exploit this vulnerability by undergoing bioreductive activation selectively in hypoxic regions, generating cytotoxic radical species that induce DNA double-strand breaks and prevent repopulation of therapy-resistant hypoxic cell compartments. Tirapazamine (TPZ), the prototypical member of the 1,2,4-benzotriazine 1-oxide HAP class^22,23^, demonstrated hypoxia-selective cytotoxicity and clinical promise but ultimately failed in Phase III trials^24^, motivating extensive analogue development efforts. Subsequent HAPs such as SN30000 exhibited improved extravascular diffusion and potency in preclinical spheroid and xenograft models when combined with gemcitabine, with the key finding that hypoxic cells initially spared by chemotherapy were effectively eliminated when SN30000 was administered concurrently or prior to chemotherapy exposure^25–28^. Despite their therapeutic potential, many HAP and TPZ analogue candidates have encountered clinical development barriers related to unfavorable pharmacokinetic profiles and dose-limiting toxicities. However, HAP evofosfamide demonstrated hypoxia-selective activity in osteosarcoma models and, when combined with Dox *in vivo*, produced enhanced tumor suppression whilst notably reducing chemotherapy-induced bone destruction^29^, highlighting the therapeutic potential of HAP and chemotherapy combinations to improve efficacy and mitigate treatment-associated morbidity. Yet despite these advances, the structure-activity relationships governing HAP potency, particularly regarding modifications to the benzotriazine core structure, remain incompletely understood. There is a need to characterize the endogenous molecular responses that HAPs induce in 3D tumor models, leveraging spatial molecular imaging to map direct treatment-induced biological adaptations and inform how HAPs can be rationally developed and optimally combined with conventional chemotherapy.

Here we investigate TPZ-A-X, a tirapazamine analogue bearing a saturated cyclopentane ring fused to the aromatic heterocycle while retaining the amino group at the 3-position of the triazole ring, representing a structural configuration that has not been previously examined for biological activity. We evaluate TPZ-A-X in combination with Dox across treatment-naïve and newly developed DXR osteosarcoma MCTS models, revealing cell-line-specific synergistic and antagonistic dose responses that necessitated deeper mechanistic investigation into how prior treatment history and microenvironmental context influence combination drug efficacy.

To determine the molecular features that govern how microenvironmental constraints and acquired resistance phenotypes influence HAP/chemotherapy combination efficacy, we applied integrated spatial multimodal MSI approaches. Imaging Mass Cytometry (IMC) was employed to map single-cell level protein expression across metabolic, survival signaling, cell cycle, DNA damage, and extracellular matrix markers. Desorption Electrospray Ionization MSI (DESI-MSI) enabled untargeted spatial metabolomics to uncover metabolic pathway enrichment and identify treatment induced metabolic reprogramming. Laser Ablation-Inductively Coupled Plasma MSI (LA-ICP-MSI) investigated endogenous metal redistribution, which serve as indicators of DNA repair capacity and apoptotic regulation. Matrix-Assisted Laser Desorption/Ionization MSI (MALDI-MSI) assessed cellular energy balance and redox state to distinguish adaptive metabolic reprogramming from cytotoxic stress. And critically, by performing DESI-MSI and IMC on the same tissue culture sections, protein and metabolite data were integrated at the individual spheroid level, enabling discovery of functional protein-metabolite correlations that reveal systems-level adaptations driving drug resistance heterogeneity. This multimodal spatial omics strategy has provided comprehensive mechanistic insight into the coordinated metabolic, signaling, and microenvironmental regulation of biological responses to HAP/chemotherapy combination exposure, informing strategies for HAP development and optimization of combination treatment strategies.

## RESULTS AND DISCUSSION

### Dose responses of hypoxia-activated prodrug and doxorubicin combination treatments

Synergistic dose responses were observed following Tirapazamine analogue X (TPZ-A-X) and Dox combination treatments in Osteosarcoma SAOS-2 MCTS models (Figure 1a). In contrast, MG-63 models showed an opposing antagonistic response to the same combination treatments (Figure 1b). Treatment-naïve SAOS-2 MCTS models were analyzed for combination responses (Figure 1a), while a broader comparison across MG-63 2D monolayers and 3D MCTS, including both treatment-naïve and DXR variants, was performed (Figure 1b). Across these analyses, drug resistance was consistently higher in 3D MCTS compared with 2D cultures, with resistance further enhanced in the DXR models relative to their treatment-naïve counterparts.

**Figure 1.**
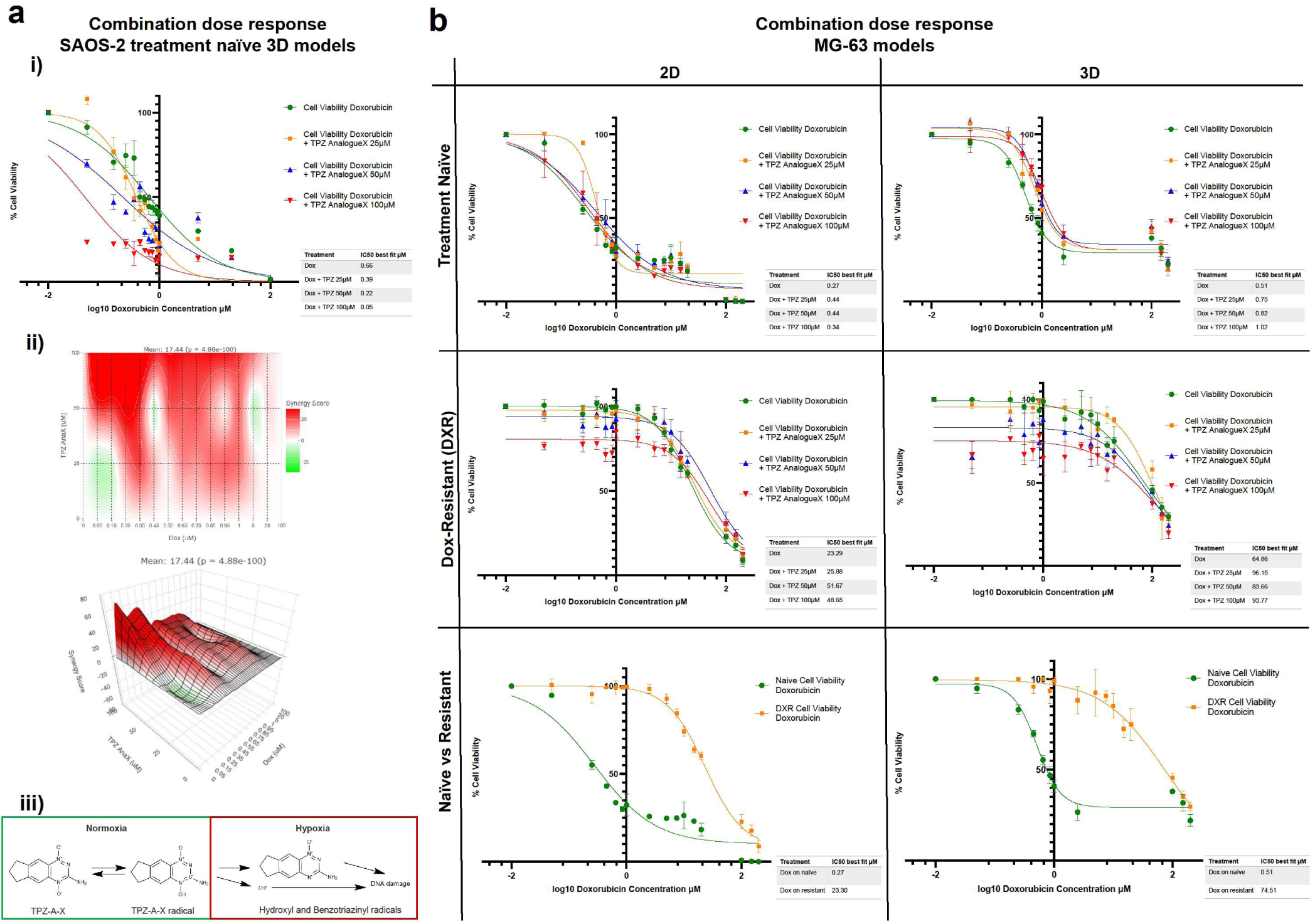
Dose responses to TPZ-A-X and Dox combination treatments (48hr) in osteosarcoma cells measured using the CellTiter-Glo 3D cell viability assay. ATP levels were determined as a function of Relative Luminescence Units (RLUs). (a) (i) Treatment-naïve SAOS-2 MCTS models treated with Dox (0.05–100 µM), alone and in combination with 25, 50, or 100µM TPZ-A-X. Data represents means +/- SD (n = 6). (ii) Synergy was quantified using an excess-over-Bliss model, and visualized as a heat map and 3D response surface, calculating a mean synergy score of 17.44 (p = 4.88 × 10⁻¹⁰⁰). (iii) A schematic of HAP mechanism of TPZ-A-X is shown. (b) MG-63 models were treated with Dox (0.05–200 µM) alone and in combination with 25, 50, or 100µM TPZ-A- X. Responses are shown across 3D MCTS, 2D monolayers, treatment-naïve, and DXR variants. Data represents means +/- SD (n = 6).

### TPZ-A-X potentiates doxorubicin-mediated cytotoxicity in SAOS-2 spheroids, producing a synergistic dose-response effect through the hypoxia-activated prodrug mechanism

The synergistic relationship observed in SAOS-2 MCTS models following combination treatment is consistent with the hypoxia-activated mechanism of TPZ-A-X, which undergoes a one-electron reduction to form an oxygen-sensitive radical anion^30^. In normoxia, the tautomers exist in a state of dynamic equilibrium. Under hypoxic conditions, the protonated intermediate yields reactive hydroxyl and benzotriazinyl radicals which can induce DNA damage (Figure 1a iii). The addition of TPZ-A-X therefore potentiates Dox mediated cytotoxicity in 3D spheroids, which are known to develop oxygen gradients, forming hypoxic regions that support HAP activation. This synergistic effect is shown in the marked left-shift of the combination dose-response curves in SAOS-2 MCTS models (Figure 1a i). The excess over Bliss synergy scoring model (Figure 1a ii) determined that an effect of 17.44% excess over expected under independent action was observed following the combination treatment with Dox, indicating the synergistic functional contribution of TPZ-A-X under conditions where hypoxia-driven activation can occur, potentially due to hypoxic cells exhibiting greater resistance to chemotherapy agents. The TPZ analogue SN30000 in combination with gemcitabine exhibited a similar response in colorectal adenocarcinoma spheroids and xenografts^26^. For Osteosarcoma, HAP combination studies with chemotherapy are limited. The TPZ analogue, evofosfamide, showed hypoxia-selective cytotoxicity in MG-63 and SAOS-2 models^31^. Combination treatments with Dox in another osteosarcoma cell line, BTK-143, studied *in vivo*, showed that combination treatment enhanced tumor suppression compared with Dox alone^31^. Synergistic *in vivo* responses led to reduced osteosarcoma-induced bone destruction with the addition of evofosfamide. Whilst this study represents the closest comparison to our investigation of TPZ-A-X and Dox, there are substantial chemical differences between evofosfamide and the TPZ analogue used here. Given these structural distinctions, the TPZ-A-X dose-response and its interaction with Dox represent a previously unexamined combination, assessed here in an unexamined human osteosarcoma model system.

### MG-63 spheroids show reduced sensitivity to doxorubicin when dosed in combination with TPZ-A-X, reflecting cell-line-specific metabolic programming and microenvironmental adaptation differences

In contrast, MG-63 models did not exhibit the synergistic interaction between TPZ-A-X and Dox. Instead, combination treatments produced an opposing trend, with IC₅₀ values for Dox remaining relatively unchanged or marginally increased with increased TPZ-A-X dose (Figure 1b). Intrinsic differences in differentiation state and osteogenic programming between cell lines may contribute to variations in hypoxic responsiveness and drug metabolism^32,33^, potentially reflecting variation in hypoxic microenvironment development and associated shifts in regulated cell-death pathways. This makes it important to investigate the factors that underpin these responses. Our previous studies of osteosarcoma ‘aggregoid’ 3D models demonstrated that MG-63 and SAOS-2 cell lines develop distinct metabolic phenotypes in 3D culture, reflecting differences in microenvironmental adaptation^17^. Furthermore, we showed that MG-63 and SAOS-2 differ markedly with respect to hypoxia markers, in that aggregoids of MG-63 express GLUT-1, a marker of hypoxia, throughout aggregoids, whereas SAOS-2 express GLUT-1 primarily in the core hypoxic regions, and not in the normoxic periphery^17^. This may suggest that the response observed here could be due to MG-63 existing in a state of pseudohypoxia, with elevated HIF-responses, which can induce heme-oxygenase-1, lowering oxidative stress. Linked to both Dox and TPZ-A-X, HIF can drive SLC11A7/xCT expression, facilitating cystine import crucial for glutathione synthesis and protection from reactive oxygen species (ROS), inhibiting both TPZ-A-X actions and those reported to drive Dox-resistance in 3D osteosarcoma cell cultures^16^. Furthermore, tirapazamine-dominant combinations have been reported to produce antagonism, while Dox-dominant ratios generated synergy^34^. Although the antagonism mechanism is not directly investigated, this reflects the trend observed here in MG-63 models, where TPZ-A-X did not enhance Dox efficacy at the analyzed dose ratios.

### Three-dimensional spheroid culture amplifies drug resistance in MG-63 models, with acquired doxorubicin-resistance further increasing resistance, revealing microenvironment-dependent resistance mechanisms

In MG-63 models, the defining feature of the response profile was the significant increase in drug resistance in 3D MCTS compared with 2D cultures, with resistance further increased in the DXR models relative to treatment-naïve counterparts (Figure 1b). Collectively, the MG-63 dose response curves (Figure 1b) highlight a model in which drug resistance is driven by microenvironmental context and prior treatment history. The clear separation between 2D and 3D cultures, and between treatment-naïve and DXR variants, provides an informative basis for investigating the mechanisms that underlie combination drug responses. MG-63 models were therefore prioritized for further analysis to determine the molecular features that contribute to how microenvironmental constraints and acquired resistance phenotypes influence the efficacy of HAP/chemotherapy combinations.

### Single-cell level spatial mapping of protein expression by Imaging Mass Cytometry (IMC)

Analysis across the four treatment groups revealed marked shifts in protein expression between DXR models compared with treatment-naïve counterparts, and in models dosed with the HAP/Dox combination treatment compared with the untreated (Figure 2). Proteins showed differential regulation, with some influenced primarily by drug resistance, others by acute treatment, and some by both factors in combination. Notably, more heterogeneity of protein expression between individual spheroid models was observed in the DXR tumor models than in the treatment-naïve. The four treatment groups (treatment-naïve untreated (control), treatment-naïve acutely treated with TPZ-A-X 50μM/Dox 0.15μM, DXR untreated (control) and DXR acutely treated with TPZ-A-X 50μM/Dox 0.15μM) each consisted of n=15 spheroids (Figure 2). Shifts in spatial localization and remodeling in response to treatment and resistance state were observed, shown in Figure 2a, with single-channel protein images presented in Figure S2, and the percentage of positive cells for each marker, together with the distribution of weak, moderate, and strong expression, is presented in Figure 2b–i, characterizing intratumoral spatial remodeling in response to treatment and resistance state.

**Figure 2.**
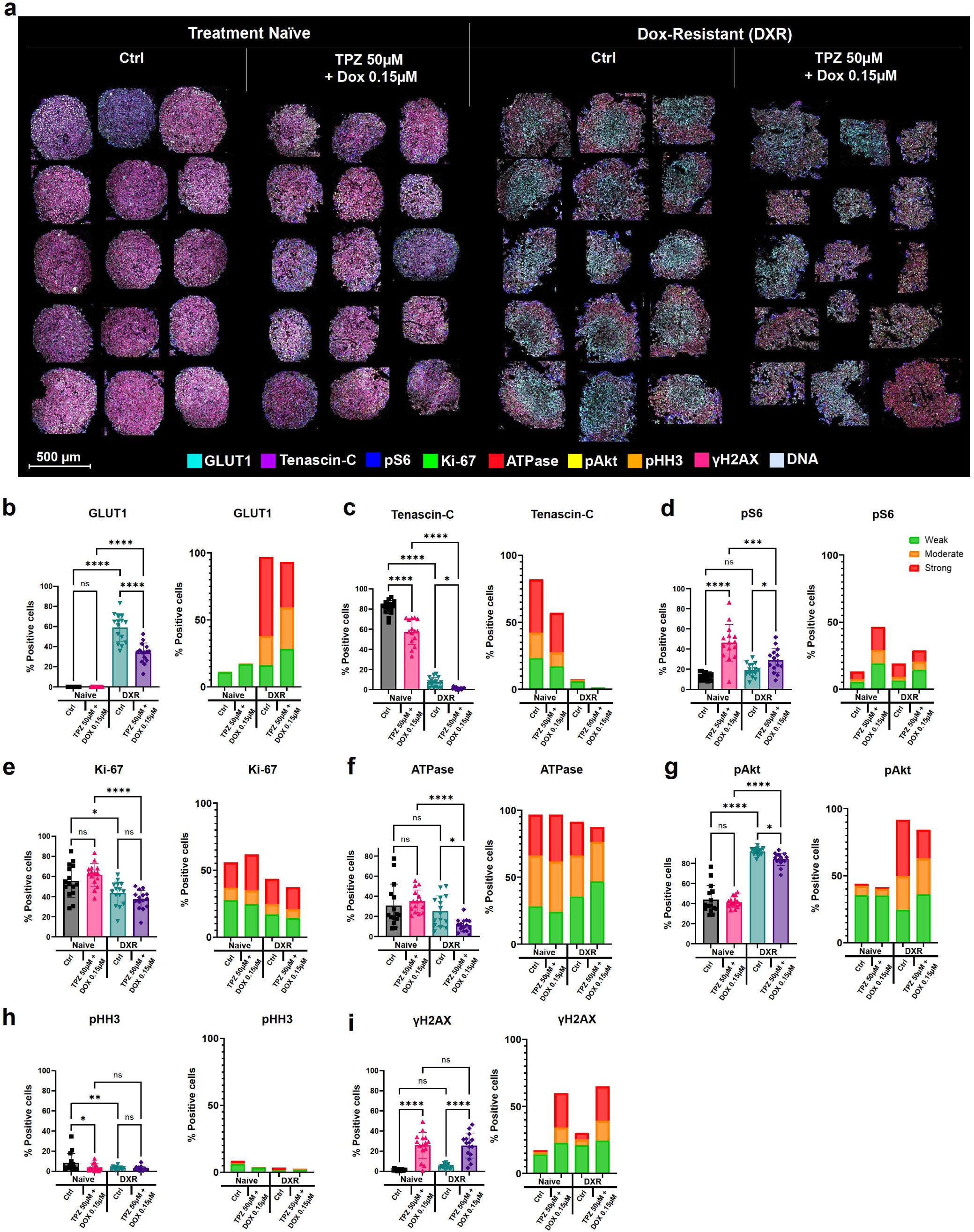
Spatial Protein analysis of MG-63 MCTS models across four treatment groups; treatment-naïve untreated (control), treatment-naïve acutely treated with TPZ-A-X 50μM/Dox 0.15μM, DXR untreated (control) and DXR acutely treated with TPZ-A-X 50μM/Dox 0.15μM. n=15 spheroids per group. (a) Protein localization analyzed at 1μm spatial resolution by IMC, shown by overlayed images of GLUT1 (cyan), Tenascin-C (purple), pS6 (blue), Ki-67 (green), ATPase (red), pAkt (yellow), pHH3 (orange), γH2AX (pink) and DNA (grey). (b) Percentage of positive cells are plotted for each spheroid model (n=15), in addition to stacked bars showing % of positive cells with average protein intensity categorized as weak, moderate or strong expression. Comparisons are made across treatment groups for the detection of GLUT1; (c) Tenascin-C; (d) pS6; (e) Ki-67; (f) ATPase; (g) pAkt; (h) pHH3; (i) and γH2AX. Significance: ns P ≥ 0.05; * P < 0.05; ** P < 0.01; *** P < 0.001; **** P < 0.0001.

### Therapeutic pressure increases protein expression heterogeneity across tumor models

Varying levels of heterogeneity were observed across individual spheroids within each treatment group, with the treatment-naïve control group exhibiting noticeably homogenous protein expression compared with the treated or resistant groups (Figure 2). The increased regional variation in these latter populations may reflect the broader range of adaptive behaviors that emerge under therapeutic pressure, whereby cells diverge into distinct subpopulations with different adaptive states of protein regulation. Heterogeneity was further assessed and is presented in Figure 4 and Figure S6.

**Figure 3.**
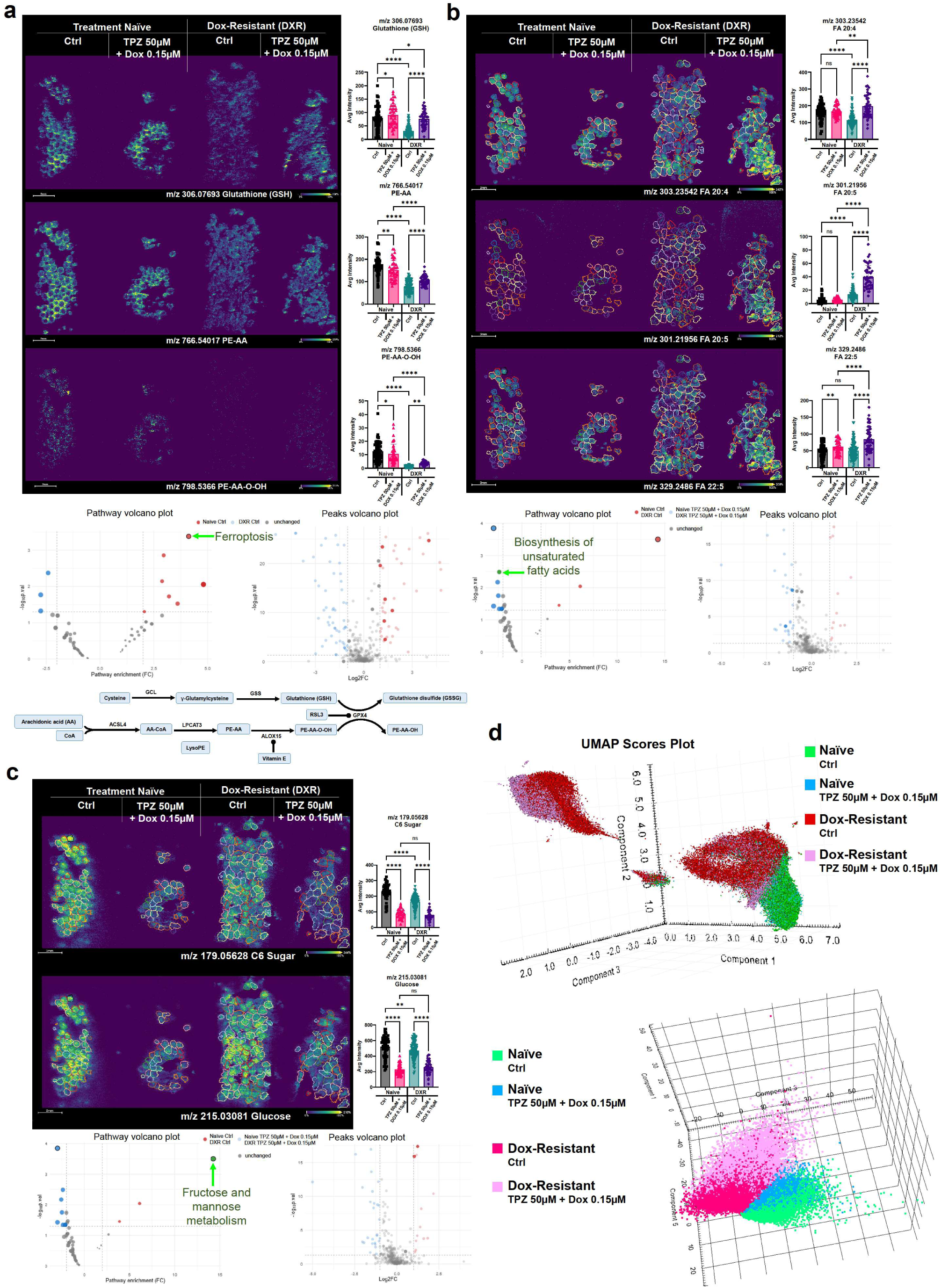
Metabolic pathway enrichment analysis of MG-63 MCTS models across four treatment groups; treatment-naïve untreated (control), treatment-naïve acutely treated with TPZ-A-X 50μM/Dox 0.15μM, DXR untreated (control) and DXR acutely treated with TPZ-A-X 50μM/Dox 0.15μM. Molecular imaging carried out by DESI-MSI at 30μm spatial resolution. Significance: ns P ≥ 0.05; * P < 0.05; ** P < 0.01; *** P < 0.001; **** P < 0.0001. (a) Ferroptosis pathway enrichment comparing untreated DXR to treatment-naïve controls. Ion density maps displayed, with corresponding average intensities plotted for; GSH (m/z 306.07693); PE-AA (m/z 766.54017) and PE-AA-O-OH (m/z 798.5366). The pathway volcano plot and peaks volcano plot show the pathway enrichment fold change against p values, naïve models (orange) and DXR models (blue). (b) Biosynthesis of unsaturated fatty acids pathway enrichment comparing acute combination treatment (TPZ-A-X 50μM/Dox 0.15μM) to untreated controls across both naïve and resistant groups. Ion density maps displayed, with corresponding average intensities plotted for; FA 20:4, e.g., arachidonic acid (m/z 303.23542); FA 20:5, e.g., eicosapentaenoic acid (m/z 301.21956) and FA 22:5, e.g., docosapentaenoic acid (m/z 329.2486). The individual tumor model segmented region annotations are shown. The volcano plots compare untreated (orange) and combination dosed (blue) models. (c) Fructose and mannose metabolism pathway enrichment comparing acute combination treatment to untreated controls across both naïve and resistant groups. Ion density maps displayed, with corresponding average intensities plotted for; C6 sugar (m/z 179.05628) and glucose (m/z 215.03081). The volcano plots compare untreated (orange) and combination dosed (blue) models. (d) Unsupervised MVA demonstrated resistance-dependent and acute dose-dependent separation. The UMAP and PCA scores plot points are color coded by treatment group: treatment-naïve untreated (green), treatment-naïve TPZ-A-X 50μM/Dox 0.15μM dosed (blue), DXR untreated (red/dark pink) and DXR TPZ-A-X 50μM/Dox 0.15μM dosed (pink).

**Figure 4.**
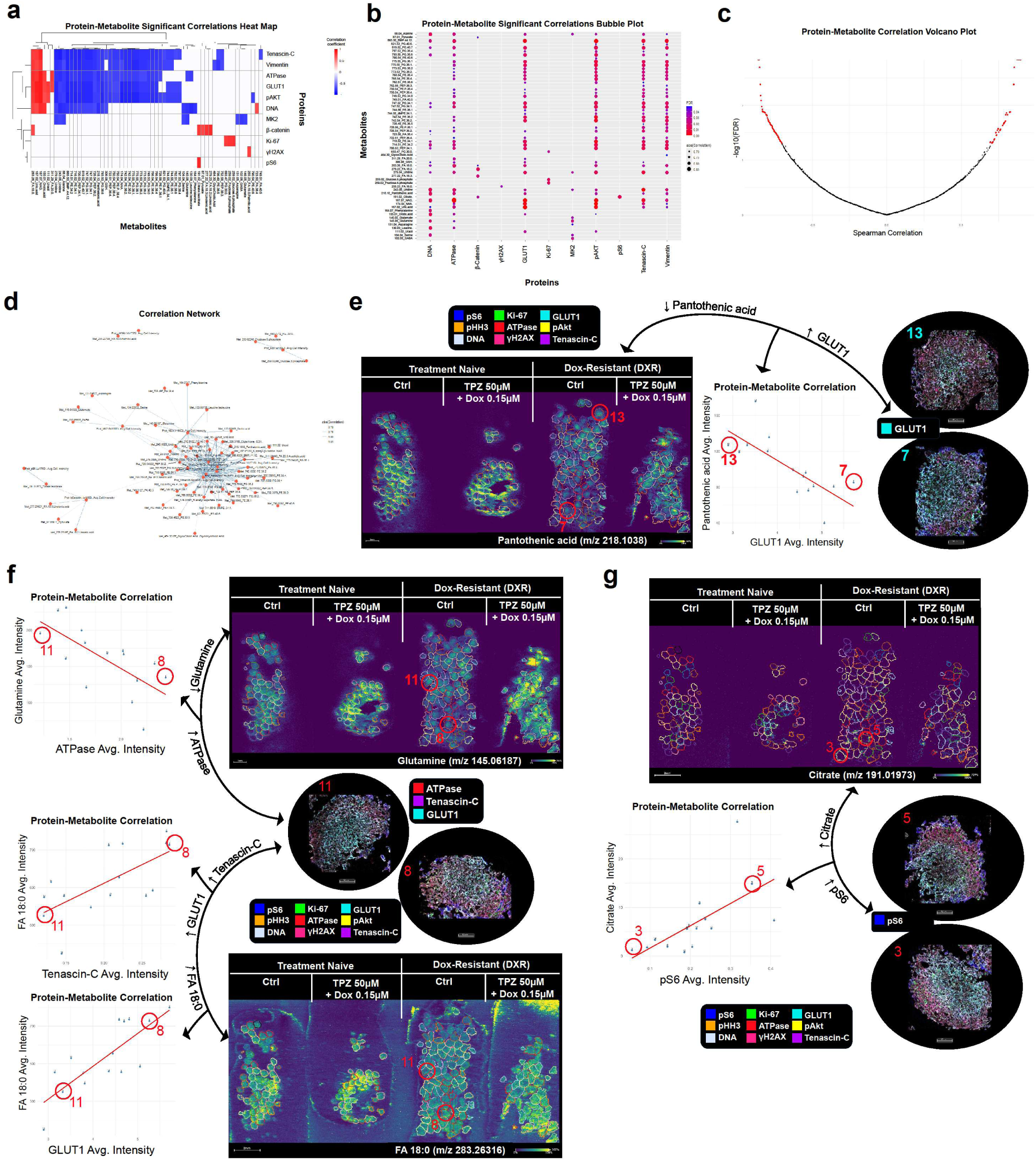
Correlative protein-metabolite analysis integrating DESI-MSI and IMC data from MG-63 MCTS models within the DXR treatment group. (a) Heatmap displaying protein–metabolite correlation coefficients. Metabolites (x-axis), proteins (y-axis), and the color scale indicating correlation strength and direction (red = positive correlation, blue = negative correlation). (b) Bubble plot visualizing protein-metabolite correlations, whereby size represents the absolute correlation value across proteins (x-axis) and metabolites (y-axis), and FDR-adjusted p-values represented by the color scale. (c) Volcano plot showing Spearman’s correlation coefficients against the –log(FDR-adjusted p-value) for protein–metabolite pairs. Significant correlations (FDR < 0.05) are highlighted in red, denoting top hit relationships. (d) Protein–metabolite correlation network demonstrating interconnected relationships within DXR tumor models. Nodes represent proteins or metabolites and edges indicate significant correlations (Spearman, FDR < 0.05). Network topology is quantified through degree centrality and edge density, highlighting clusters, hubs, and key molecules underlying functional connectivity. (e–g) Correlation plots of selected protein–metabolite relationships. Each data point represents an individual spheroid model, plotting metabolite intensity against protein intensity. (e) Negative correlation between pantothenic acid abundance and GLUT1 expression, visualized alongside representative IMC and DESI ion images from spheroid models #13 and #7. (f) Negative correlation between glutamine abundance and ATPase expression (top), and a positive correlation between FA 18:0 abundance and Tenascin-C and GLUT1 expression (bottom). (g) Positive correlation between citrate abundance and pS6 expression, co-localized to the outer tumor edge. n = 15 individual spheroid models correlated.

### Doxorubicin-resistance induces distinct microenvironmental and signaling adaptations

The most striking observations in DXR spheroids were the potent induction of GLUT1 coupled with increased pAkt, and a pronounced decrease in Tenascin-C (TNC) detection. (Figure 2b). Spatially, within each individual tumor model, GLUT1 expression was largely homogenous within the regions of treatment-naïve spheroids, whereas DXR models, particularly untreated, showed increased core localization of GLUT1 (Figure 2a and Figure S2), suggesting that the enhanced glycolytic capacity supporting acquired resistance here is spatially enriched within the hypoxic core regions of resistant spheroids. The core localization of GLUT1 positions the glycolytic machinery where oxygen availability is most limited, effectively coupling metabolic adaptation to the hypoxic microenvironment that drives chemoresistance. The upregulation of GLUT1 in resistant models reflects a shift toward glycolytic metabolism, enabling enhanced glucose uptake to sustain ATP production and redox balance under stress; a metabolic adaptation foundationally understood as a hallmark of cancer but also more recently associated with chemoresistance and poor prognosis in various cancer types, including osteosarcoma^35^. In Dox-resistant osteosarcoma cells specifically, this glycolytic shift has been mechanistically linked to upregulation of glucose transporters and glycolytic enzymes through the ELK1/PTBP1 signaling axis^36^. The GLUT1 upregulation observed here in DXR spheroids is therefore displaying an acquired resistance phenotype which is driven by enhanced glycolytic capacity. This metabolic shift is consistent with models in which tumors reduce glycolytic dependence and increase mitochondrial ATP generation when glycolytic flux is constrained, altering their susceptibility to therapy^37,38^.

pAkt was significantly higher in DXR spheroids than in naïve spheroids, consistent with a role of PI3K/Akt activation in sustaining survival and chemoresistance (Figure 2g). The PI3K/Akt pathway blocks apoptosis by phosphorylating and inhibiting pro-apoptotic proteins such as Bcl-2-associated death promoter (Bad) and caspase-9 while upregulating anti-apoptotic factors, and it is a recognized mediator of multidrug resistance across cancer types^39–41^. In addition, Akt signaling integrates with other pro-survival pathways, including NF-κB and MAPK/ERK cascades^42^, to protect tumor cells from chemotherapy-induced stress and to promote multidrug resistance. In response to Dox, activation of upstream HER3–PI3K–Akt signaling has been shown to protect tumor cells from drug-induced stress, contributing directly to reduced apoptosis^43^. The elevated pAkt signal in DXR spheroids is therefore consistent with an acquired survival mechanism that helps maintain viability under Dox treatment.

TNC was significantly downregulated in DXR spheroids compared with naïve controls (Figure 2c). This was unexpected given that TNC is widely described as a pro-tumorigenic extracellular matrix (ECM) glycoprotein that promotes survival, invasion, and resistance to apoptosis in several solid tumors, in part through modulation of integrin, EGFR, and NF-κB signaling^44–46^. High TNC expression within the tumor microenvironment has also been associated with cancer stemness and therapy resistance by supporting niche-like ECM architectures^47^. Loss of TNC in the DXR spheroids may reflect a differentiation shift induced by acquisition of resistance. TNC is associated with a mesenchymal phenotype, and its reduction is consistent with the looser spheroid formation observed in DXR spheroids, likely mediated by decreased ECM production and altered differentiation. Consistent with this, TNC silencing has been reported to decrease osteogenic markers, including Col1A1 and alkaline phosphatase, in osteoblast-like cells, supporting a model in which TNC levels modulate differentiation rather than simply mark it in osteosarcoma^48^.

### Hypoxia-activated prodrug/doxorubicin combination treatment differentially alters metabolic, survival and ECM protein expression in naïve and resistant spheroids

The TPZ-A-X/Dox combination treatment further altered these protein expression patterns in a manner that differentiated the acute response observed in naïve and resistant spheroids. TNC showed a further decrease following TPZ-A-X/Dox combination treatment in both naïve and DXR models. This further decrease may reflect treatment-induced remodeling of the spheroid microenvironment away from a TNC-rich matrix state (Figure 2c). Given that TNC has been linked to the maintenance of cancer stem cell niches in other tumor types^49,50^, its reduced expression in untreated DXR spheroids suggests that, in this osteosarcoma model, acquired Dox-resistance does not require a TNC-rich, stem-like microenvironment. The further reduction of TNC following TPZ-A-X/Dox treatment in both naïve and DXR spheroids may reflect preferential cytotoxicity towards cells that are more dependent on TNC-rich ECM support, leaving behind populations that maintain viability through TNC-independent mechanisms (Figure 2c). While speculative, this may indicate a model in which TNC-associated niches preferentially support more therapy-responsive cells, whereas long-term resistant cells rely less on this ECM component. Overall, these data suggest that within the spheroid microenvironment, both acquired Dox-resistance and the acute response to HAP/Dox treatment occur without dependence on TNC-mediated ECM support, distinguishing this resistance mechanism from TNC-driven chemoresistance described in other cancers. Also key to the response observed here was the spatial distribution remodeling which TNC underwent across treatment groups. In treatment-naïve untreated models, TNC expression was homogenously distributed throughout the whole spheroid region, but following acute TPZ-A-X/Dox combination treatment there was a loss of peripheral expression with a shift toward core localization (Figure 2a and Figure S2), indicating acute drug induced ECM remodeling away from the periphery. In contrast, untreated DXR spheroids displayed preferential outer-edge localization of TNC, possibly reflecting enrichment of a subset of resistant cells that maintain limited TNC dependency at the spheroid periphery. This spatial redistribution pattern aligns with the proposed model whereby TNC-rich niches are progressively lost through selective cytotoxicity to sensitive cells, whereas resistant populations persist through TNC-independent mechanisms. These intratumoral localization shifts further support the idea that resistance in this model is independent of TNC-mediated ECM support.

HAP/Dox combination treatment of treatment-naïve and DXR spheroids also revealed marked increased detection of pS6 and γH2AX in tumor models following the acute exposure, irrespective of resistance status (Figure 2d, i). The increased percentage of pS6-positive cells following acute treatment (Figure 2d) supports the trends observed in our previous work, where pS6 was identified as a marker of acute Dox non-responding cells^14^. Phosphorylated ribosomal protein S6 is a downstream readout of mTORC1 activity and reflects activation of anabolic, growth-promoting signaling pathways that enhance protein synthesis, cell growth, and nutrient sensing^51^. High pS6 levels have been associated with poor therapeutic response and earlier recurrence in several cancers, and reduced S6 phosphorylation has been linked to improved outcome under targeted therapy^52,53^. The absence of baseline differences between naïve and DXR models indicates that pS6 is not a defining feature of the acquired resistance phenotype in the osteosarcoma spheroid models. Instead, the selective increase in pS6 following acute combination dosing suggests that TPZ-A-X/Dox combination treatment triggers an adaptive mTOR response that is engaged in reaction to drug exposure, particularly in naïve models where the upregulation is more pronounced than in DXR spheroids (Figure 2d). This supports pS6 as a candidate biomarker of acute treatment adaptation to the HAP/Dox therapy, rather than of long-term resistance, and crucially implies that mTOR-dependent signaling is initiated to buffer the stress associated with combination-induced DNA damage and metabolic disruption. Spatially, pS6 expression was predominantly localized to the outer edge of spheroids in all treatment groups, but showed depth-dependent redistribution following acute combination dosing (Figure 2a and Figure S2). In untreated models, pS6 was tightly localized to the peripheral edge, but following TPZ-A-X/Dox exposure, pS6-positive cells remain mostly in the periphery but are extended deeper into the spheroid, supporting the idea of a subpopulation of mTOR-active, drug-tolerant cells which survive the combination dose and expand beyond their initial peripheral niche. In the context of the strong γH2AX induction (Figure 2i), pS6 upregulation may reflect an attempt by surviving cells to support repair and survival pathways. This raises interest for the investigation of combination treatments with mTOR inhibitors. In addition, the increased γH2AX following treatment is consistent with stalled replication forks in response to topoisomerase inhibition by Dox, observed even in DXR models. Along with the coexistence of strong γH2AX and elevated pAkt in the resistant spheroids (Figure 2g, 2i), this also suggests that although the combination treatment successfully induces substantial DNA damage, the acquired survival mechanism, dampens the ability of these lesions to drive full apoptotic commitment.

A significant reduction in GLUT1 expression was also observed following combination treatment, specifically in DXR models with no change observed in naïve models (Figure 2b). This selective effect suggests that the combination therapy disrupts the metabolic reprogramming that underpins Dox-resistance. The reduction in GLUT1 may force a shift toward oxidative phosphorylation, potentially sensitizing resistant cells to subsequent cytotoxicity following the combination dose. Decreased glycolysis and increased oxidative phosphorylation following the combination dose to DXR tumor models may have facilitated the dose response observed in Figure 1b, which lead to more ATP output and subsequent shift in dose response. Glycolysis has previously been directly implicated in osteosarcoma chemoresistance^54^, with glycolytic inhibition by 2-deoxy-D-glucose significantly enhancing the antitumor efficacy of Dox in osteosarcoma xenograft models^55^. In the response shown here, the selective reduction of GLUT1 in DXR spheroids following TPZ-A-X/Dox combination treatment suggests that the combination disrupts the glycolytic adaptation that supports acquired resistance, providing new understanding, which builds on previous glycolysis-chemoresistance links, to the glycolytic inhibition capabilities of the TPZ-A-X/Dox combination treatment (Figure 2b). This indicates that the combination treatment targets a metabolic dependency that is more pronounced in resistant than in naïve populations. The downregulation of glycolytic markers in response to HAP treatment, and specifically in combination with Dox, has not been reported previously in osteosarcoma, and has not been characterized in 3D MCTSs of any cancer type. Collectively, these findings reveal a new mechanistic insight by which HAP/Dox combination treatment selectively reduces GLUT1 expression in DXR tumor models but not in treatment-naïve models, implying a metabolic reprogramming of glycolytic dependence as a key component to the HAP/Dox sensitization revealed in a hypoxia-structured microenvironment.

### Untargeted metabolomic pathway enrichment analysis reveals distinct metabolic signatures of acquired resistance and acute hypoxia-activated prodrug and doxorubicin combination responses

To complement the spatial protein analysis, untargeted metabolomic imaging by DESI-MSI was performed on the same slide of tumor models, investigating the four treatment groups with each treatment group consisting of ∼40-100 tumor models, n=64, 41, 98, 47 respectively (Figure 3). Pathway enrichment analysis was carried out to identify metabolic pathways that were disproportionately represented among differentially expressed metabolites, highlighting the key drivers of metabolic divergence between treatment groups. Enrichment across multiple metabolic pathways was revealed, with several providing key understanding to both acquired Dox-resistance and acute combination treatment responses, which are presented here (Figure 3). Specifically, pathway enrichment identified suppression of ferroptosis-associated pathways as a hallmark of the DXR phenotype, alongside distinct treatment-induced signatures in unsaturated fatty acid biosynthesis and fructose/mannose metabolism. These metabolic adaptations align with the protein-level inhibition of glycolytic flux and suggest a coordinated reprogramming of energy metabolism and redox homeostasis. MVA further supported the findings that show the acute combination treatment induces a metabolic state distinct from both naïve and stable resistant phenotypes, indicating a unique therapeutic stress signature (Figure 3). Molecular annotations and corresponding mass errors are detailed in Figure S4. Additional pathway enrichment results including all top pathway hits are presented in Figure S3.

### Ferroptosis markers are suppressed in doxorubicin-resistant cells, but reversed by TPZ-A-X/doxorubicin combination treatment

Pathway enrichment analysis comparing untreated DXR models to treatment-naïve controls identified the ferroptosis pathway as a key discriminator of the DXR phenotype (Figure 3a). Ferroptosis is a regulated cell death cascade triggered by iron-dependent accumulation of lipid peroxides to lethal levels, resulting in oxidative membrane damage and cell death^56^. A schematic of the ferroptosis cascade is shown in figure 3a, outlining the glutathione biosynthesis and arachidonate metabolism portion of the pathway. The decrease of ferroptosis-associated metabolites in DXR spheroids reveals a striking adaptation: glutathione (GSH) (m/z 306.07693), phosphatidylethanolamine-arachidonic acid (PE-AA) (m/z 766.54017), and its oxidized hydroperoxide form (PE-AA-O-OH) (m/z 798.5366) were all significantly depleted relative to naïve controls (Figure 3a).

GSH is a critical cofactor for glutathione peroxidase 4 (GPX4), the primary enzyme responsible for neutralizing lipid peroxides and preventing ferroptotic cell death^57^. The reduced baseline GSH in DXR spheroids might initially suggest metabolic vulnerability; however, in the context of acquired chemoresistance, this depletion likely reflects regulation of redox homeostasis or a shift toward alternative antioxidant pathways that limit oxidative stress without triggering death. More critically, the decrease of PE-AA and PE-AA-O-OH in resistant spheroids indicates suppression of the lipid substrate pool. PE-AA is a primary target for oxidation by lipoxygenases (LOXs) to generate lethal lipid peroxides that initiate ferroptosis^58^. By maintaining lower levels of these oxidizable lipids, DXR cells may raise the threshold for ferroptotic triggering. Therefore, employing a mechanism of ferroptosis evasion, increasingly recognized as a strategy employed by drug resistant cells^59–64^ (Figure 3a).

Critically, acute combination treatment with TPZ-A-X/Dox reversed this ferroptosis signature. In DXR models, combination dosing partially restored GSH, PE-AA, and PE-AA-O-OH levels, with GSH returning almost to naïve control levels (Figure 3a). This metabolic reversal suggests that the combination treatment actively re-engages the ferroptotic machinery that resistant cells had deliberately suppressed. By replenishing the pool of oxidizable lipids and restoring GPX4 cofactor availability, the combination treatment may overwhelm the resistant cells’ redox defences, converting the previously supressed ferroptotic pathway back into an active vulnerability.

Tirapazamine has been shown to induce ferroptosis in OS through SLC7A11/GPX4 downregulation^65,66^, and ferroptosis induction is a known strategy to overcome chemoresistance^67–69^. Our findings provide novel evidence that the TPZ-A-X/Dox combination specifically reverses the adaptive ferroptosis suppression characterized by GSH and lipid depletion in DXR spheroids. The restoration of ferroptosis capacity observed may therefore represent a critical component of how TPZ-A-X and Dox synergistically overcome acquired resistance (Figure 3a).

### Upregulation of unsaturated fatty acids reflects lipid peroxidation substrate accumulation

In parallel with the restoration of ferroptosis markers, pathway enrichment analysis identified a significant increase of unsaturated fatty acid biosynthesis following acute combination treatment (Figure 3b). Metabolites of the enriched biosynthesis of unsaturated fatty acids pathway included increased FA 20:4, e.g., arachidonic acid (m/z 303.23542), FA 20:5, e.g., eicosapentaenoic acid (m/z 301.21956) and FA 22:5, e.g., docosapentaenoic acid (m/z 329.2486) in combination-dosed models. Notably, FA 20:4 and FA 20:5 showed a significant increase specifically in DXR combination-dosed spheroids but not in naïve combination-dosed models, whereas FA 22:5 increased in both groups, with a more pronounced increase in DXR models (Figure 3b).

This treatment-induced accumulation of polyunsaturated fatty acids (PUFAs) mechanistically links to the re-engagement of ferroptosis described above in Figure 3a. PUFAs such as arachidonic acid are the essential substrates for lipoxygenase-catalysed lipid peroxidation; their incorporation into membrane phospholipids directly sensitizes cells to ferroptotic collapse^58,70,71^. This may reflect a two-fold strategy: the combination treatment not only restores ferroptotic capacity, via GSH and PE-AA/O-OH restoration (Figure 3a), but also supplies the substrate, elevated PUFAs, needed for ferroptosis (Figure 3b). Alternatively, this may represent a compensatory membrane remodelling response^72,73^, where cells attempt to repair HAP-induced oxidative damage to their lipid bilayers but inadvertently increase their susceptibility to further peroxidation.

### Universal glycolytic substrate depletion and inhibition of glycolytic flux

In contrast to the resistance-specific lipid remodelling, pathway enrichment for fructose and mannose metabolism revealed a universal response to combination treatment across both naïve and DXR populations (Figure 3c). Key glycolytic entry substrates, including hexose sugars (m/z 179.05628) and glucose (m/z 215.03081), were significantly decreased in combination dosed models, regardless of resistance status (Figure 3c). This finding is supported by the GLUT1 downregulation observed in Figure 2b, which strongly supporting a model of treatment induced glycolytic flux inhibition. This may represent a fundamental effect of the combination treatment and affects both naive and DXR populations.

### Spatial and multivariate metabolic discrimination reveals resistance-dependent and treatment-dependent grouping

Unsupervised MVA showed that the metabolic states defined by resistance status and acute treatment are spatially and biochemically distinct (Figure 3d). PCA scores plots demonstrated clear discrimination between treatment-naïve and DXR models, validating that acquired resistance is accompanied by a fundamental, stable shift in the global metabolomic phenotype. Acutely treated models also separated from their untreated counterparts within each group, confirming that the drug-induced metabolic responses, such as those presented here showing the ferroptosis restoration (Figure 3a), PUFA accumulation (Figure 3b), and hexose depletion (Figure 3c), represent significant departures from baseline homeostasis.

Spatially, UMAP analysis revealed component specific localization patterns with important implications for understanding how the HAP/Dox combination engages the spheroid microenvironment (Figure 3d, Figure S5). Component 1, driven by metabolic features characteristic of naïve models, was also present in DXR spheroids but restricted to the outer, more oxygenated periphery of the resistant models. Conversely, Components 2 and 3, representing DXR specific metabolic features, were predominantly enriched in the spheroid core. This spatial distribution highlights the influence of the hypoxic gradient on metabolic phenotype. The presence of naïve like metabolic signatures localised to the peripheral zone of resistant spheroids suggests that the core, where oxygen is most limited and TPZ-A-X bioreduction and radical generation are maximal, represents a metabolically distinct region which is dominated by resistance associated adaptations.

This spatial organization is mechanistically of interest because it reveals that the HAP/Dox combination is targeting the metabolic vulnerabilities precisely where resistance mechanisms are most firmly established (Figure 3d). The hypoxic core, where resistance-specific metabolic features are shown to be concentrated (Components 2 and 3), is also the region where signatures of ferroptosis suppression is the greatest (low GSH, low PE-AA/OOH) and where glycolytic dependence is highest (baseline GLUT1 upregulation). By simultaneously acting in this core region to restore ferroptotic substrates (increased GSH, PE-AA and PUFAs) while depleting glycolytic fuels (decreased glucose and hexoses), the combination therapy may create a multi-directional metabolic collapse specifically in the compartment where resistant cells are most vulnerable and least able to adapt.

### Integrative protein-metabolite correlation analysis reveals functional links between metabolic reprogramming and signaling adaptations driving drug resistance heterogeneity

Integrating multimodal DESI-MSI and IMC data from individual tumor models enabled investigation of protein-metabolite relationships within the same tissue-culture section, revealing functional links that may underpin drug resistance mechanisms, differential treatment response, and intragroup heterogeneity in spheroid populations (Figure 4). Protein-metabolite correlation analysis was used to identify significant relationships within the treatment group examined, DXR MG-63 MCTS models. By leveraging the spatial resolution of these imaging data, we identified protein-metabolite correlations and quantified the extent to which inter-spheroid heterogeneity within a single treatment group contributed to these relationships (Figure 4 and Figure S6). Deviation from group level trends, defined by z scores, were used to calculate a heterogeneity index (H). The resulting distributions reveal marked spheroid-to-spheroid variability across protein–metabolite pairs (Figure S6).

### Identifying key protein–metabolite correlations and network connectivity within drug-resistant models

Proteins and metabolites interact closely, and their interplay forms an integral part of the complex systems that drive disease progression and influence how cells respond or fail to respond to treatments^74^. The heatmap visualizations shown in Figures 4a and 4bguided the selection of key relationships of interest subsequently presented in Figures 4e-g. The volcano plot presented in Figure 4c further guided the identification of significant protein-metabolite pairs by plotting Spearman correlation coefficients against –log(p-values). Through this integrated analysis, relationships highlighted in red (Figure 4c), were identified as the most significant correlations, FDR < 0.05. These were subsequently mapped into a correlation network presented in Figure 4d. The greater interconnectivity of the pantothenic acid–GLUT1 pair within the larger network hub implies that glucose transport and CoA-related metabolism form part of a shared regulatory network maintaining cellular energy balance. While the lower interconnectivity of the citrate–pS6 pair points to a more spatially distinct, niche-driven signaling event (Figure 4d).

### Individual spheroid heterogeneity in protein–metabolite relationship adaptations revealed through integrated multimodal spatial analysis

The examination of individual spheroid models within the DXR cohort revealed substantial heterogeneity in protein-metabolite relationships (Figure S6). This approach uncovered how protein-metabolite associations vary across different spheroids, allowing the identification of the most informative bivariate relationships displayed in Figure 4e–g.

Figure 4e demonstrates a significant negative correlation between pantothenic acid abundance and GLUT1 expression. Spheroid models with elevated pantothenic acid levels displayed reduced GLUT1 expression, whereas spheroids with diminished pantothenic acid showed compensatory increases in GLUT1. This relationship is exemplified by comparing spheroid #13 and #7: spheroid #13 exhibits lower GLUT1 expression, indicated by IMC, and correspondingly higher pantothenic acid abundance, evident in the DESI-MSI ion images, whereas spheroid #7 shows the opposing pattern (Figure 4e). While the direct regulatory link between pantothenic acid availability and GLUT1 expression demonstrated here (Figure 4e) has not been previously described, this correlation can be attributed to compromised CoA biosynthesis when pantothenic acid is limited ^75^, resulting in cells not sustaining efficient oxidative phosphorylation and compensating via the upregulation of glucose import via GLUT1 ^76^ to meet energetic demands through glycolysis.

A significant negative correlation between ATPase expression and glutamine abundance was observed in Figure 4f. Tumor models with lower glutamine levels had significantly elevated ATPase expression, whereas spheroids maintaining higher glutamine levels showed reduced ATPase activity. This inverse relationship indicates metabolic reprogramming in response to glutamine availability. ATPase enzymes, particularly ATP synthase and other ATP-consuming processes, represent substantial energy expenditure. Glutamine fuels ATP generation through oxidative phosphorylation via glutaminolysis^77^; when it becomes limiting in the tumor microenvironment, cells face reduced bioenergetic capacity^78^. The elevated ATPase expression observed in glutamine-scarce spheroids suggests a compensatory mechanism where enhanced ATP synthesis is required to sustain energy-demanding cellular processes, such as DNA repair and protein synthesis, that can no longer be adequately supported by glutamine-derived ATP^79,80^.

Positive correlations between FA 18:0, e.g., stearic acid abundance and both TNC and GLUT1 expression were observed (Figure 4f). Spheroids with elevated stearic acid levels showed increased expression of both TNC and GLUT1, suggesting that lipid-enriched regions of the tumor microenvironment, characterized by abundant saturated fatty acids, coincide with enhanced glucose uptake capacity and lipid driven extracellular matrix remodeling. Stearic acid, particularly when incorporated into membrane structures, influences membrane biophysics and can modulate protein trafficking and signaling through lipid modification^81,82^.

A significant positive correlation between citrate abundance and pS6 expression was observed in Figure 4g, with the distinctive feature that the metabolite and protein exhibit co-localization at the outer edge of DXR spheroid models (Figure 4g). Citrate and pS6 have not been previously linked at the single-spheroid level, although citrate is a key substrate for mTORC1-driven anabolism and provides acetyl-CoA for biosynthetic pathways, and so similarly to pS6 ^83,84^, it is associated with mTOR driven resistance. High pS6 levels where citrate accumulates may suggests robust mTORC1-dependent anabolism. This anabolic state at the spheroid periphery demonstrated here, can be considered alongside the treatment group comparative IMC analysis previously discussed in Figure 2d, to further understand the responses observed whereby pS6 marked acute treatment non-responsive, proliferative cells. Here, the peripheral concentration of both citrate and pS6 reflects the favorable metabolic microenvironment: oxygen-rich, nutrient-rich conditions proximal to the culture medium enabling robust oxidative phosphorylation and mTORC1-driven growth signaling. The inner hypoxic regions experience suppressed mTORC1 signaling and reduced biosynthetic capacity, consistent with oxygen-dependent nutrient sensing^85^. Collectively, this integrative interpretation, enabled by multimodal spatial correlation, reveals a coordinated systems-level adaptation and suggests functional integration influenced by protein and metabolite heterogeneity across individual spheroid models.

### Cellular redox state and energy balance assessment reveals intact metabolic viability despite endogenous metal depletion in hypoxia-activated prodrug/doxorubicin combination treated models

LA-ICP-MSI was performed to map endogenous metal localization, while MALDI-MSI was performed to map endogenous metabolites and subsequently assess cellular redox state and energy balance across the same treatment groups. By the integration of these complementary imaging modalities, metal and metabolite imaging, multi-dimensional evidence showed a depletion of endogenous metal abundance in combination-treated DXR models, and importantly, this occurs independently of oxidative stress or energy dysregulation, supporting a tumor model of functionally viable metabolic reprogramming rather than general cellular toxicity (Figure 5). Strikingly, this metal redistribution is accompanied by significant spatial reorganization of copper and magnesium, revealing selective cofactor redeployment that aligns with the spheroid microenvironment and sites of optimal HAP activation.

**Figure 5.**
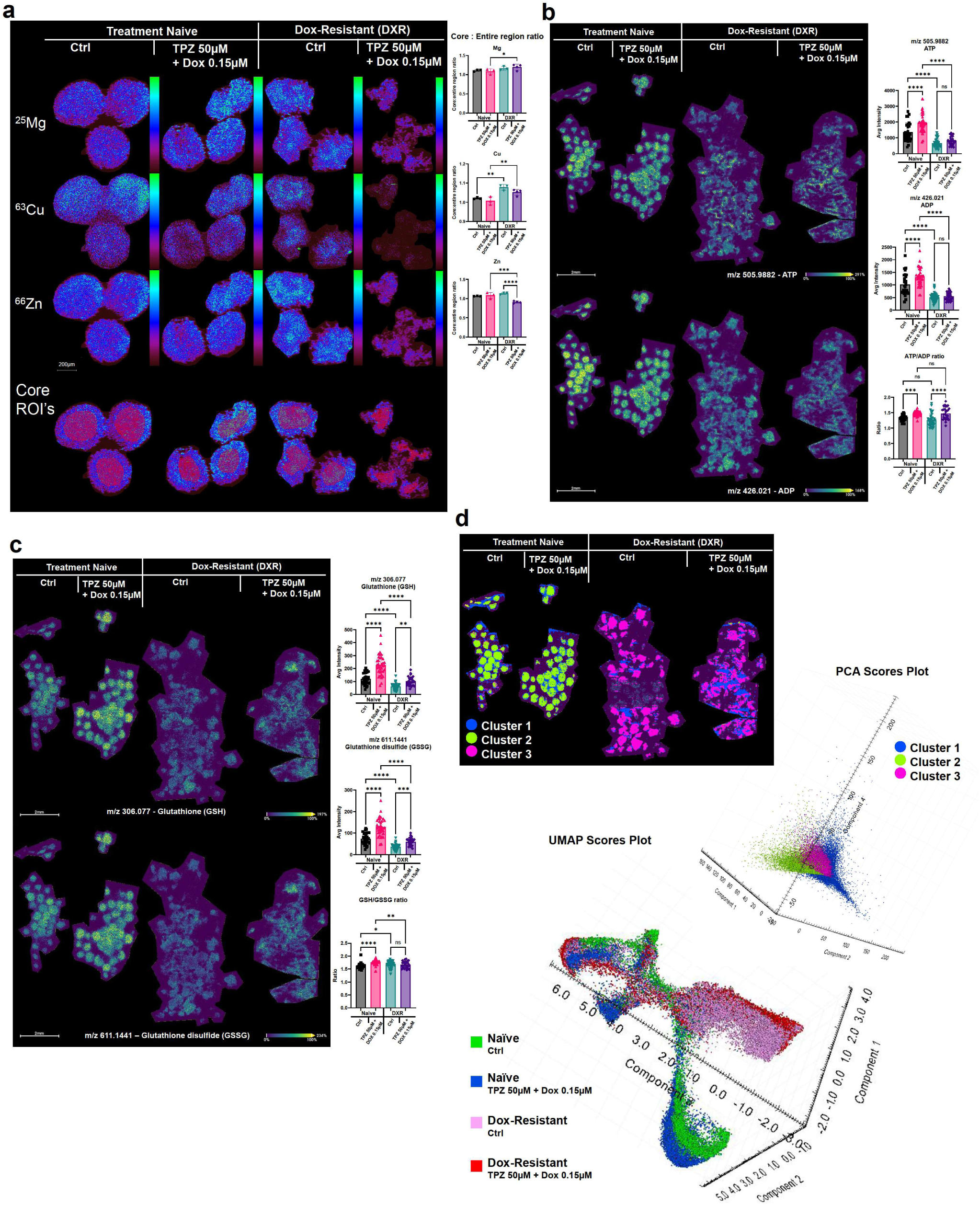
Endogenous metal abundance, cellular energy balance, and redox state analysis across MG-63 MCTS models across four treatment groups; treatment-naïve untreated control, treatment-naïve acutely treated with TPZ-A-X 50μM/Dox 0.15μM, DXR untreated control, and DXR acutely treated with TPZ-A-X 50μM/Dox 0.15μM. (a) Endogenous metal analysis by LA-ICP-MSI at 5μm spatial resolution. Ion images of magnesium (^25^Mg), copper (^63^Cu), and zinc (^66^Zn) are displayed across treatment groups and their corresponding core:entire region ratios are shown. Core region of interest (ROI) images used for ratio calculations are shown. Within each treatment group, n=3 spheroids were analyzed. (b-c) MALDI-MSI analysis to investigate (b) cellular energy balance by ATP/ADP ratio shown alongside ion density maps of ATP (m/z 505.9882) and ADP (m/z 426.021), with corresponding average intensities plotted, and (c) cellular redox state and oxidative stress by glutathione/glutathione disulfide (GSH/GSSG) ratio shown alongside ion density maps of GSH (m/z 306.077) and GSSG (m/z 611.1441), with corresponding average intensities plotted. Within each treatment group, 29-42 spheroids were analyzed (n = 30, 36, 42, 29 across the treatment groups left to right). Significance: ns P ≥ 0.05; * P < 0.05; ** P < 0.01; *** P < 0.001; **** P < 0.0001. (d) Unsupervised MVA of MALDI-MSI spectra. PCA was conducted, and spatial segmentation was performed using PCA component scores. The PCA scores plot and spatially mapped clusters are shown. Three clusters were identified: cluster 2 (green), dominated by treatment-naïve spheroids; cluster 3 (pink), corresponding to DXR models; and cluster 1 (blue), localized to spheroid peripheries and extracellular regions shared by both conditions. UMAP analysis was also performed and the UMAP scores plot revealed metabolic separation by; component 1 dominated by DXR regions, component 2 by naïve models, and component 3 spanning both resistance states.

### Endogenous metal redistribution occurred in response to acute hypoxia-activated prodrug/doxorubicin exposure

LA-ICP-MSI identified distinct shifts in endogenous metal localization across treatment groups. Among magnesium and zinc, copper exhibited the most striking spatial redistribution (Figure 5a). In treatment-naïve spheroids, copper showed relatively homogeneous distribution throughout the tumor model, with a slight shift to the outer edge away from the tumor core following treatment. In contrast, DXR models had pronounced core-localized copper accumulation, with core:entire region ratios significantly elevated (Figure 5a). This spatial redistribution was specific to the resistant phenotype across both untreated and acutely combination dosed (Figure 5a).

Copper’s role as a catalytic cofactor in redox and DNA repair enzymes^86–89^ provides mechanistic insight into this adaptation. High intracellular copper concentration enhances the activity of copper-dependent oxidases and facilitates rapid DNA damage recognition and repair^90,91^, potentially contributing to enhanced resistance capacity in DXR models. The pronounced core-localized copper accumulation in untreated DXR spheroids may reflect an acquired resistance strategy (Figure 5a). In addition, recent reports have linked copper accumulation to the regulation of cuproptosis, a newly characterized form of regulated cell death distinct from apoptosis and ferroptosis^92,93^. Given that copper accumulation itself can trigger cuproptosis when levels exceed cellular regulatory capacity, the core-localized enrichment observed here raises an intriguing therapeutic question as to whether targeting this adaptive copper accumulation could convert it from a resistance mechanism into a vulnerability. Cuproptosis requires activated TCA cycle flux and lipoylated (post-translational modification with lipoic acid) TCA enzymes, with key targets being DLAT (E2 subunit of pyruvate dehydrogenase) and DLST (alpha-ketoglutarate dehydrogenase complex)^92,93^, meaning that copper becomes toxic when it can engage the TCA machinery. Since HAPs consume GSH and oxidized thiols, and GSH and metallothioneins bind Cu and Zn^94,95^, in the presence of GSH decrease, metal ions can be exported e.g. via ATP7A/B^96^, consistent with the observations shown here (Figure 5a and 5c).

### ATP/ADP ratio indicates maintained bioenergetic capacity despite profound magnesium depletion

Magnesium showed a treatment-dependent, resistance-specific pattern (Figure 5a). In treatment-naïve models, acute combination treatment produced an increase in magnesium intensity. In contrast, DXR models showed a marked decrease in magnesium following combination dosing (Figure 5a and Figure S7). MALDI-MSI analysis of ATP and ADP demonstrated that cellular energy balance is preserved in HAP/Dox treated DXR spheroids despite the marked reduction in intracellular magnesium. When ATP and ADP were assessed individually, both ATP and ADP increased following acute combination treatment in naïve models but remained unchanged in DXR spheroids (Figure 5b). This divergence suggests that naïve cells actively expand their adenine nucleotide pools to meet the elevated energetic demands imposed by drug-induced stress, whereas resistant cells maintain stable, albeit lower, baseline ATP and ADP levels. Importantly, the ATP/ADP ratio increased in HAP/Dox treated spheroids, indicating that adenylate energy charge remains intact^97^. Given that dying or metabolically collapsing cells typically exhibit a sharp fall in ATP, a rise in ADP, corresponding to a decrease in ATP/ADP ratio, the observed increase provides strong evidence that these spheroids remain bioenergetically competent ^98,99^. Thus, the stability of ATP, ADP, and their ratio confirms that the pronounced magnesium depletion reflects a regulated metalomic adaptation rather than nonspecific cytotoxicity or loss of viability (Figure 5b). This reduction in glycolytic flux ^100–102^ following the HAP/Dox combination dose aligns with the observations of GLUT1 downregulation, (Figure 2b), and the hexose depletion (Figure 3c). Together, these findings support a model in which the combination treatment forces a metabolic shift: cells restrict glucose uptake and glycolytic throughput, thereby reducing ATP demand and maintaining sufficient energy reserves through modulated oxidative phosphorylation ^103^.

Magnesium homeostasis is dynamically regulated through both intracellular sequestration and transmembrane transport, and hypoxia-induced stresses are known to perturb these pathways^104^. One plausible explanation is that HAP activation within hypoxic spheroid cores disrupts mitochondrial Mg²⁺ retention. Mitochondrial Mg²⁺ uptake, primarily mediated by the Mrs2 channel^105^, is sensitive to membrane potential; hypoxia-driven reductions to membrane potential would impair Mg²⁺ uptake and promote its release into the cytosol and ultimately the extracellular space. Additionally, several Mg²⁺ exporters, most notably SLC41A1 and the Na⁺/Mg²⁺ exchanger, are activated under oxidative and metabolic stress. Upregulation of SLC41A1 has been linked to chemoresistance and metabolic rewiring in other systems^106^, suggesting that resistant spheroids may actively extrude Mg²⁺ as part of a stress-response program. TRPM7, a Mg²⁺-permeable channel with kinase activity, is also responsive to intracellular ROS and ATP levels^107^. Altered TRPM7 activity under dual treatment could further contribute to a net loss of cellular Mg²⁺. These transport-linked mechanisms are regulated processes, not passive leakage, and the maintenance of ATP despite Mg²⁺ depletion indicates that the Mg loss represents a targeted metabolic adaptation to combined HAP/Dox exposure rather than a consequence of cell death or membrane failure. This work highlights a novel mechanism of profound magnesium depletion without accompanying ATP/ADP ratio collapse under drug stress.

Zinc showed a significant reduction in core:entire region ratio in DXR models following combination treatment (Figure 5a, S7 and S8), indicating redistribution away from the hypoxic core toward the periphery. Zinc serves as a structural and catalytic cofactor for numerous DNA repair enzymes, including zinc-finger proteins and DNA polymerases^108,109^. The peripheral redistribution of zinc following combination treatment may reflect increased demand for zinc-dependent repair machinery in the more oxygenated, peripheral tumor regions where cells are attempting to manage treatment-induced damage while maintaining proliferative capacity (Figure 5a). Alternatively, zinc depletion from the core may represent zinc mobilization to support ATP-dependent processes or antioxidant defense in response to acute oxidative stress^109^. However, coupled with Cu and Mg observations, these data support a model in which hypoxia-activated drug metabolism triggers a broad disruption of metal homeostasis, indicative of a true metallome collapse, that is mechanistically decoupled from adenylate energy balance, representing a previously unrecognised cellular stress phenotype.

### Preserved GSH/GSSG ratio confirms absence of oxidative stress despite hypoxia-activated prodrug/doxorubicin combination treatment

The glutathione redox couple (GSH/GSSG) serves as an indicator of cellular oxidative stress and redox homeostasis^110,111^. Elevated GSSG relative to GSH, indicated by a decreased GSH/GSSG ratio, reflects an oxidatively stressed state characterized by increased lipid and protein damage^112^. Notably, the GSH/GSSG ratio following acute combination treatment observed here contradicts this expected stressed state decrease. An increase following acute treatment was observed in treatment-naïve, and maintenance in DXR models (Figure 5c). This ratio maintenance is mechanistically significant. Individual GSH and GSSG absolute intensities increased following HAP/Dox treatment in both Naïve and DXR, reflecting increased metabolic flux through the glutathione biosynthesis pathway^113^, whilst the stoichiometric ratio crucially remained intact. This indicates that the combination treatment does not generate uncontrolled oxidative stress or overwhelm cellular antioxidant defenses. Instead, the proportional increase in both oxidized and reduced glutathione suggests a coordinated upregulation of redox metabolism rather than a stress-induced redox imbalance^114^ (Figure 5c).

This preserved redox homeostasis is particularly striking given that HAPs such as TPZ-A-X are designed to generate reactive oxygen and nitrogen species under hypoxic conditions^115,116^. The fact that GSH/GSSG ratios remain stable or increased suggests that the glutathione system successfully buffers the oxidative challenge imposed by HAP-derived radicals, preventing lethal oxidative damage while still allowing DNA damage to accumulate, as supported by γH2AX regulation and discussed in Figure 2i. The simultaneous loss of Cu and Zn (Figure 5a), key cofactors for metallothioneins and Cu/Zn-SOD, would further increase reliance on the glutathione system, accelerating depletion of total glutathione pools even as redox state remains controlled. This finding provides critical reassurance that the depleted metal abundances (Figure 5a) and altered energy balance observed in combination-treated spheroids (Figure 5c) occur independently of general cellular toxicity or oxidative damage. The cells maintain their capacity to regulate intracellular redox state, supporting the model that metal depletion and energy remodeling are functional, adaptive responses to drug exposure rather than markers of cellular failure.

### Multivariate analysis reveals distinct yet spatially overlapping metabolic landscapes across treatment and resistance states

PCA of MALDI-MSI spectra, combined with spatial segmentation using the PCA component scores as the feature data, identified three distinct metabolic clusters across treatment groups (Figure 5d). The PCA scores plot is color coded by these identified clusters which are separated by PCA components 1, 2 and 4. As shown on the spatial segmentation map, Cluster 2 (green) is dominated by treatment-naïve spheroid regions and represents a treatment naïve metabolic signature. Cluster 3 (pink) corresponds to DXR models and reflects the hypoxic core metabolic phenotype characterized by resistance-associated adaptations. Cluster 1 (blue) is localized to spheroid peripheries and extracellular regions shared by both treatment-naïve and resistant states, representing a transitional metabolic state (Figure 5d). This spatial organization reflects the dominant influence of both the spheroid microenvironment and acquired resistance-associated metabolic reprogramming on metabolic phenotype: the aerobic, nutrient-rich periphery supports metabolic states partially shared between naïve and resistant models, while the hypoxic core exhibits resistance-specific metabolic and elemental adaptations driven by the metabolic phenotype acquired through prolonged Dox exposure.

### Spatial UMAP analysis further stratified metabolic complexity in a resistance and treatment-dependent manner

UMAP Component 1, driven by DXR characteristic metabolic features, dominates the spheroid core where hypoxia is most pronounced and TPZ-A-X bioactivation is maximal (Figure 5d and Figure S8). This aligns with the core-localized copper accumulation and metal redistribution observed in Figure 5a. UMAP Component 2 is driven by treatment-naïve metabolic signatures and is distributed throughout naïve spheroids but also present in the periphery of DXR models, suggesting metabolic plasticity at the tumor edge. Component 3, spanning both resistance states and concentrated in extracellular and peripheral regions which are accessible to nutrient and oxygen diffusion, has identified a shared peripheral metabolic signature accessible to both naïve and resistant phenotypes (Figure 5d and Figure S8). This layered metabolic stratification, where DXR specific signatures concentrate in the core, treatment-naïve signatures dominate throughout naïve spheroids, and a shared peripheral metabolic signature bridges both resistance states in the nutrient-rich outer regions, demonstrates that the spheroid microenvironment and acquired resistance phenotype together enforce distinct yet spatially overlapping metabolic identities.

## CONCLUSIONS

Understanding how microenvironmental context and acquired resistance phenotypes influence the efficacy of combination therapies remains a critical challenge in anticancer drug development. This study applied integrated spatial multimodal MSI to map the coordinated metabolic, signaling, and microenvironmental responses that characterize how HAP and chemotherapy combinations can overcome Dox-resistance in osteosarcoma spheroid models.

A central finding is that TPZ-A-X/Dox combination treatment elicits a spatially distinct response in the hypoxic core, where resistance-associated metabolic adaptations are most pronounced. The combination simultaneously disrupts multiple interconnected survival pathways: it reverses ferroptosis suppression through restoration of glutathione and lipid peroxide substrates, restricts glycolytic flux via GLUT1 downregulation and hexose depletion, and suppresses pro-survival signaling through pAkt reduction. Critically, these adaptations are selective for resistant cells, with minimal effects on treatment-naïve counterparts, indicating that the therapy specifically exploits the metabolic dependencies that resistant cells have acquired.

The spatial organization of these responses reveals multi-layered mechanistic complexity. Copper and magnesium redistribution concentrate in the hypoxic core, establishing a constrained DNA repair microenvironment whilst simultaneously suppressing apoptotic escape pathways. Protein-metabolite correlations demonstrate functional coupling: glucose import capacity (GLUT1) balances against CoA-dependent oxidative metabolism (pantothenic acid), lipid composition correlates with nutrient uptake and ECM remodeling, and anabolic signaling (pS6) concentrates in nutrient-rich peripheries where citrate supports biosynthesis. These coordinated adaptations are not uniformly distributed and differ spatially according to microenvironmental demand, enabling resistant cells to maintain functional viability. Preserved ATP/ADP and GSH/GSSG ratios confirm intact bioenergetics and redox buffering yet maintain a metabolic state in which defensive resources are steadily depleted through active repair and survival signaling. This revealed, under drug-induced stress, a novel mechanism of profound magnesium depletion without an accompanying ATP/ADP ratio collapse. Together, these findings suggest that the combination does not indiscriminately kill resistant cells but instead engages their metabolic vulnerabilities by simultaneously restoring ferroptotic capacity, depleting glycolytic substrates, and weakening survival barriers, creating multi-directional metabolic failure, where no single adaptation can counter the induced response. The heterogeneity observed between individual tumor models within resistant populations points towards the value of combination therapies which target metabolic adaptations and circumvent adaptive escape routes, highlighting the need for combinations that overwhelm rather than merely inhibit compensatory mechanisms.

These findings have implications for rational HAP development and combination strategy optimization. First, the selective targeting of resistant phenotypes suggests that HAPs may offer particular clinical value in tumors that have developed chemotherapy-resistance through metabolic reprogramming. Second, the specificity of ferroptosis re-engagement and glycolytic disruption in the hypoxic core suggests why conventional chemotherapy alone shows limited efficacy in resistant tumors. Third, the multimodal integration of spatial proteomics, metabolomics and endogenous metal analysis demonstrates an approach that enables discovery of functional relationships that are inaccessible to single-modality approaches.

Collectively, these findings establish a framework for understanding how spatial metabolic organization within tumor microenvironments contributes to therapeutic vulnerability, informing the design of more effective combination strategies for chemotherapy-resistant osteosarcoma and potentially other solid malignancies.

## MATERIALS AND METHODS

### Tirapazamine analogue X (TPZ-A-X) Synthesis

Synthesis of TPZ-A-X was carried out *via* minor amendments to procedures reported by Hay et al^117,118^, over a 5-step process shown in scheme 1.

**Scheme 1.**
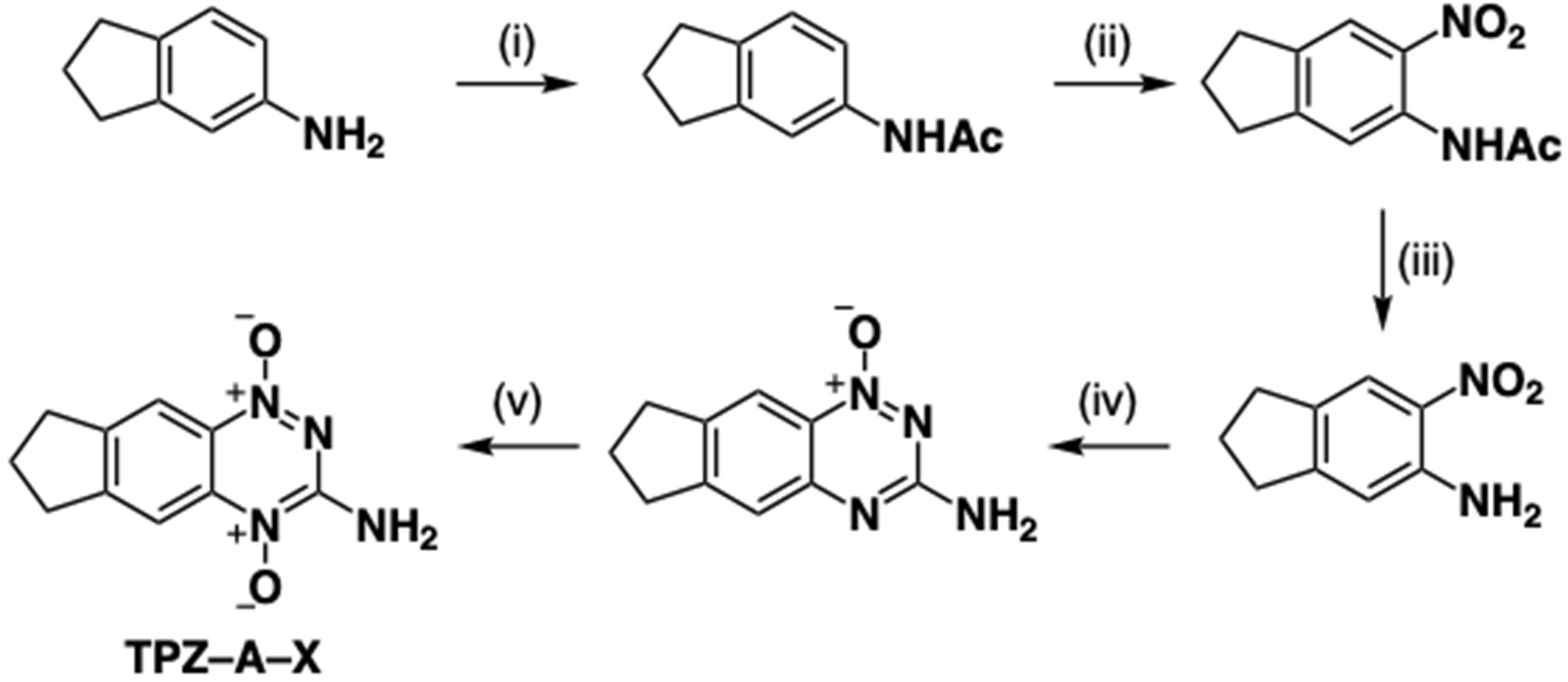
Synthesis of TPZ-A-X. (i) Ac_2_O, 1,4-dioxane, RT, 24 h; (ii) HNO_3_, AcOH, RT, 4 h; (iii) HCl (6M aq.), 100 °C, 26 h; (iv) 1. NC–NH_2_, 80 °C, 0.1 h, 2. HCl (conc.), 50–100 °C, 4 h, 3. NaOH (7.5M aq.), 50–100 °C, 16 h; (v) H_2_O_2_ (30% aq.), AcOH, 50 °C, 4 h.

### 3D cell culture tumor model formation

The human osteosarcoma cell line, MG-63 (ATCC, Manasas, VA, USA), was cultured in Minimum Essential Medium Eagle-α modification (α-MEM) growth medium, supplemented with 10% fetal bovine serum and 1% penicillin–streptomycin (ThermoFisher Scientific, Waltham, MA, USA). Cells were maintained at 37 °C, 5% CO_2_. Dox-resistant (DXR) MG-63 cells were formed by long term, stepwise increased sub-lethal doses of Dox until continued growth was achieved in constant exposure to 1μM Dox for a minimum of 40 weeks. This approach was guided by the original studies which generated resistant osteosarcoma cell lines^18^. 3D cell cultured tumor models were generated by the aggregation, from a single cell suspension, of 1 × 10^4^ cells per well in low-adhesion U-bottom plates (Greiner Bio-One Ltd., Gloucestershire, UK) to form MCTSs. MCTS models were left to form over 48hr, prior to dosing, after which the medium was replenished and treatment groups were exposed to drug-containing medium for a further 48hr, giving a total culture duration of 96hr before harvesting. Dosing media was prepared from drug stock solutions of; doxorubicin hydrochloride (Merck KGaA, Darmstadt, Germany) dissolved in water, and Tirapazamine analogue X (TPZ-A-X) dissolved in water and DMSO (highest concentration working solution <0.11% DMSO).

### Cell viability assays

CellTiter-Glo 3D Cell Viability Assays (Promega, Madison, WI, US) were carried out following treatments. The Glo-Reagent was added at a ratio of 1:1 and spheroids were incubated according to the manufacturer’s protocol^119^. Relative Luminescence Units (RLUs) were read by the ClarioStar Microplate Reader (BMG Labtech, Baden-Württemberg, Germany) as a function of the ATP concentration. Media–drug solutions in the absence of cells were employed as negative controls, from which the background was subtracted to calculate normalized ATP read outs. Cell viability was calculated and plotted in a non-linear regression fit dose response curve using GraphPad Prism version 10.4.1 (GraphPad Prism, Boston, MA, USA). Best fit IC50 values were calculated and presented. Combination dose synergy was quantified using an excess-over-Bliss synergy scoring model via the SynergyFinder+ application^120^.

### Sample preparation for multimodal mass spectrometry imaging of tumor models

At treatment end points, MCTS models were pooled by treatment groups, n = 180, washed 3× with ice-cold phosphate-buffered saline solution and embedded (n = ∼180) in 7.5% hydroxypropyl methylcellulose (HPMC) and 2.5% polyvinylpyrrolidone (PVP) as per the optimal embedding method for multi-modal molecular imaging^121^, and snap frozen using liquid nitrogen. Embedded MCTS models were cryo-sectioned (Leica Biosystems, Nussloch, Germany) at 10 μm, thaw-mounted onto poly-L-lysine-coated microscope slides (Merck KGaA, Darmstadt, Germany) or indium tin oxide slides (VisionTek Systems Ltd., Cheshire, UK) for MALDI analyses and dried immediately with a stream of compressed air to avoid molecular delocalization. Each slide was prepared to consist of a tissue section from each of the four treatment groups, to allow for same slide molecular imaging comparisons across all treatment groups. Slide mailers were vacuum-packed and stored at −80 °C.

### Desorption Electrospray Ionization - Mass Spectrometry Imaging (DESI-MSI)

Spatial metabolomics was carried out by DESI-MSI using an Exploris 480 orbitrap mass spectrometer (Thermo Fisher Scientific, Dreieich, Germany) equipped with a DESI source which used an automated stage (Prosolia Inc., Zionsville, IN, USA) and an in-house custom-built DESI sprayer. The DESI was operated at a flow rate of 1 μL/min delivering a 95:5 v/v methanol/H_2_O solvent. Nitrogen nebulizing gas was operated at 4 bars. Images were acquired at a 30 μm spatial resolution. Negative ionization mode was operated with an MS scan range of 80–1000 m/z, achieving 70,000 FWHM resolution. ThermoRAW files were converted to centroided mzML format using MSConvertGUI with MS peak picking. Each row mzML file was converted to an image, imzML file, using an imzML convertor programme^122^. Images were processed and analyzed using SCiLS Labs Software version 2026a Premium 3D (Bruker Daltonics, Bremen, Germany). Spatial segmentation was carried out using a bisecting k-means algorithm to aid region of interest annotations. Ion mean intensities were normalized by root mean square (RMS) and presented using GraphPad Prism software version 10.4.1 (GraphPad Prism, MA, USA). Unpaired parametric t tests and ordinary one-way ANOVA comparisons were performed. P values were calculated at a 95% confidence interval and annotated as follows: ns P ≥ 0.05; * P < 0.05; ** P < 0.01; *** P < 0.001; **** P < 0.0001. Putative MS1 annotations were made, with corresponding molecular formulae and mass errors detailed in Figure S4. Over-Representation Analysis (ORA) pathway enrichment methods were carried out using peak annotations and Log2 fold change calculations, plotted against p value to generate pathway enrichment and metabolite hits volcano plots. Unsupervised multivariate analysis (MVA) was carried out by principal component analysis (PCA), which used unit variance scaling to the present scores plots, and uniform manifold approximation and projection (UMAP) was carried out using a correlation distance metric and unit variance scaling to present the scores plot, using SCiLS Labs Software.

### Imaging Mass Cytometry (IMC)

Single-cell level protein localization by IMC was carried out using the Hyperion imaging system (Standard BioTools, San Francisco, CA, USA), on the same slides that were analyzed by DESI-MSI for spatial metabolomics. The laser ablation was carried out at 1 μm spatial resolution, with metal isotope detection by inductively coupled plasma time-of-flight mass spectrometry. IMC staining of the sectioned tumor models was carried out as previously described^15,16^. Briefly, the tissues were fixed with PFA, permeabilized with a Casein + 0.1% Triton X-100 solution, and blocked in Casein, before incubation with the metal conjugated antibodies (Standard BioTools, South San Francisco, CA, USA) in a humidified chamber at 4⁰C overnight. The panel of metal tag antibody conjugations and corresponding antibody targets used are detailed in Figure S1. Image analysis was carried out using HALO software v4.1 (Indica Laboratories, Albuquerque, NM, USA). The HighPlex FL algorithm was used with thresholds optimized manually. Percentage of positive cells and protein mean intensities were presented using GraphPad Prism software version 10.4.1 (GraphPad Prism, MA, USA). Unpaired parametric t tests and ordinary one-way ANOVA comparisons were performed. P values were calculated at a 95% confidence interval and annotated as follows: ns P ≥ 0.05; * P < 0.05; ** P < 0.01; *** P < 0.001; **** P < 0.0001.

### Protein-metabolite relationship correlation analysis

Spearman’s rank correlation was used to assess protein–metabolite relationships across spheroids within each treatment group, with p-values adjusted for multiple testing via the Benjamini–Hochberg FDR. Multiple comparisons analysis guided the identification and significant protein-metabolite correlation pairs. Protein and metabolite intensities were z-score standardized across all spheroids. Spheroid-specific heterogeneity indices (H) were quantified as the absolute difference between corresponding protein and metabolite z-scores. Analyses were performed in RStudio.

### Matrix-Assisted Laser Desorption/Ionization - Mass Spectrometry Imaging (MALDI-MSI)

MALDI-MSI was carried out using a timsTOF fleX mass spectrometer (Bruker Daltonics, Bremen, Germany). Sample slides were first coated with 9-aminoacridine (9AA) matrix at 10mg/mL in 80% methanol using a TM-sprayer (HTX-Technologies, Chapel Hill, NC, USA). The matrix was spayed in 6 layers using the following sprayer parameters: 75°C; 0.08 ml/min flow rate, 6 psi nitrogen pressure; 1200 mm/min velocity; 40 mm nozzle height; 3 mm track spacing in a crisscross pattern. Matrix coated samples were analyzed in negative ionisation mode, ablated to achieve at 20μm spatial resolution using a laser repetition rate of 10kHz, 80% laser power, 750 shots per pixel, and detected by the time of flight mass analyzer scanning a mass range of 200-900 m/z. Images were processed and analyzed using SCiLS Labs Software version 2026a Premium 3D (Bruker Daltonics, Bremen, Germany). Spatial segmentation was carried out using a bisecting k-means algorithm to aid region of interest annotations. Ion mean intensities were normalized by root mean square (RMS) and presented using GraphPad Prism software version 10.4.1 (GraphPad Prism, MA, USA). Unpaired parametric t tests and ordinary one-way ANOVA comparisons were performed. P values were calculated at a 95% confidence interval and annotated as follows: ns P ≥ 0.05; * P < 0.05; ** P < 0.01; *** P < 0.001; **** P < 0.0001. Putative MS1 annotations were made, with corresponding molecular formulae and mass errors detailed in Figure S4. Unsupervised MVA of the MALDI-MSI spectra was carried out, using SCiLS Labs Software, by principal component analysis (PCA) using unit variance scaling. An enhanced segmentation workflow was also carried out by using PCA component scores as the input data to apply Leiden clustering using a Euclidean metric. Uniform manifold approximation and projection (UMAP) was also performed for spatial MVA using a correlation distance metric and unit variance scaling.

### Laser Ablation–Inductively Coupled Plasma - Mass Spectrometry Imaging (LA-ICP-MSI)

LA-ICP-MSI analysis was carried out using an imageBIO266 laser ablation unit (Elemental Scientific Lasers, LLC, Bozeman, MT, USA) coupled to a NexION 5000 ICP-MS/MS (PerkinElmer, Buckinghamshire, UK). Sample slides were ablated to achieve a spatial resolution of 5µm, using a laser repetition rate of 100Hz and a fluence of 3 Jcm^−1^, whilst operating at a 25 μm/s scan speed. The MS method was optimized for the detection of magnesium (^25^Mg), copper (^63^Cu), and zinc (^66^Zn) using dwell times per atomic mass unit of 7, 10, 8, 15ms respectively. The laser ablation and MS method were optimized from an integrated imaging perspective, whereby signal to noise ratios and limits of detection^123^ were considered to devise optimal dwell times, whilst maximizing image resolution, minimizing analysis time and avoiding spectral skew imaging artefacts^124,125^. Optimization algorithms were used to apply these principles, employing statistical framework from foundational theory^126^. Images were processed and analyzed using Iolite software (version 4.10.8) (Elemental Scientific Instruments, Omaha, NE, UK).

## ASSOCIATED CONTENT

### Data Availability

The data presented in this paper are stored in the Sheffield Hallam University Research Data Archive (SHURDA) at https://shurda.shu.ac.uk.

## Author Contributions

Conceptualization: L.M.C., N.A.C., D.P.S., M.R.C., S.M.P.; Data curation: S.M.P., L.E.F., N.A.C., L.M.C.; Formal analysis: S.M.P.; Funding acquisition: M.R.C., L.M.C.; Investigation: S.M.P.; Methodology: S.M.P., L.M.C., N.A.C., D.P.S., L.E.F., D.M.A., B.J.W., J.D.R.; Project administration: L.M.C., L.E.F.; Resources: S.M.P., L.M.C., L.E.F., G.H., R.J.A.G., D.M.A., B.J.W., J.D.R.; Software: S.M.P.; Supervision: L.M.C., N.A.C., D.P.S., M.R.C., L.E.F.; Validation: S.M.P., L.M.C., N.A.C.; Visualization: S.M.P.; Writing – original draft: S.M.P.; Writing – review & editing: S.M.P., N.A.C., L.E.F., L.M.C., D.P.S., M.R.C., D.M.A., J.D.R., G.H.

## ACKNOWLEDGMENTS

This work was funded by a Sheffield Hallam University Vice Chancellor award and Waters Corporation PhD studentship. We acknowledge Astra Zeneca for their collaboration and facilitation of S.M.P’s PhD student research placements. Colleagues at the Center for Mass Spectrometry Imaging (CSMI) and the Biomolecular Sciences Research Centre (BMRC) at Sheffield Hallam University are greatly acknowledged for their facilitation of this work.

## Supporting information

Metal conjugated antibody panel (Figure S1).

IMC protein expression and individual localization images (Figure S2). Top enriched pathway hits from volcano plots (Figure S3).

Mass annotation table (Figure S4).

UMAP components of DESI-MSI mapped onto regions (Figure S5).

Heterogeneity index, z-scores across correlations, degree of deviation from group-level trends (Figure S6).

Endogenous metal intensities in entire region (Figure S7).

MALDI UMAP components mapped onto tissue regions (Figure S8).

---

**Figure S1.**
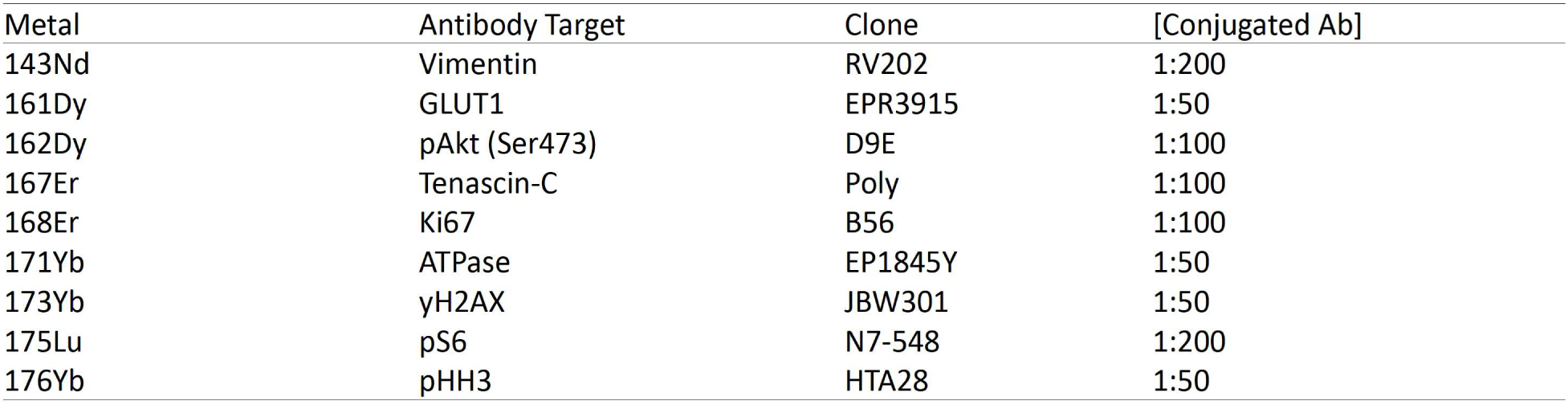
Metal conjugated antibody panel and corresponding working concentrations.

**Figure S2.**
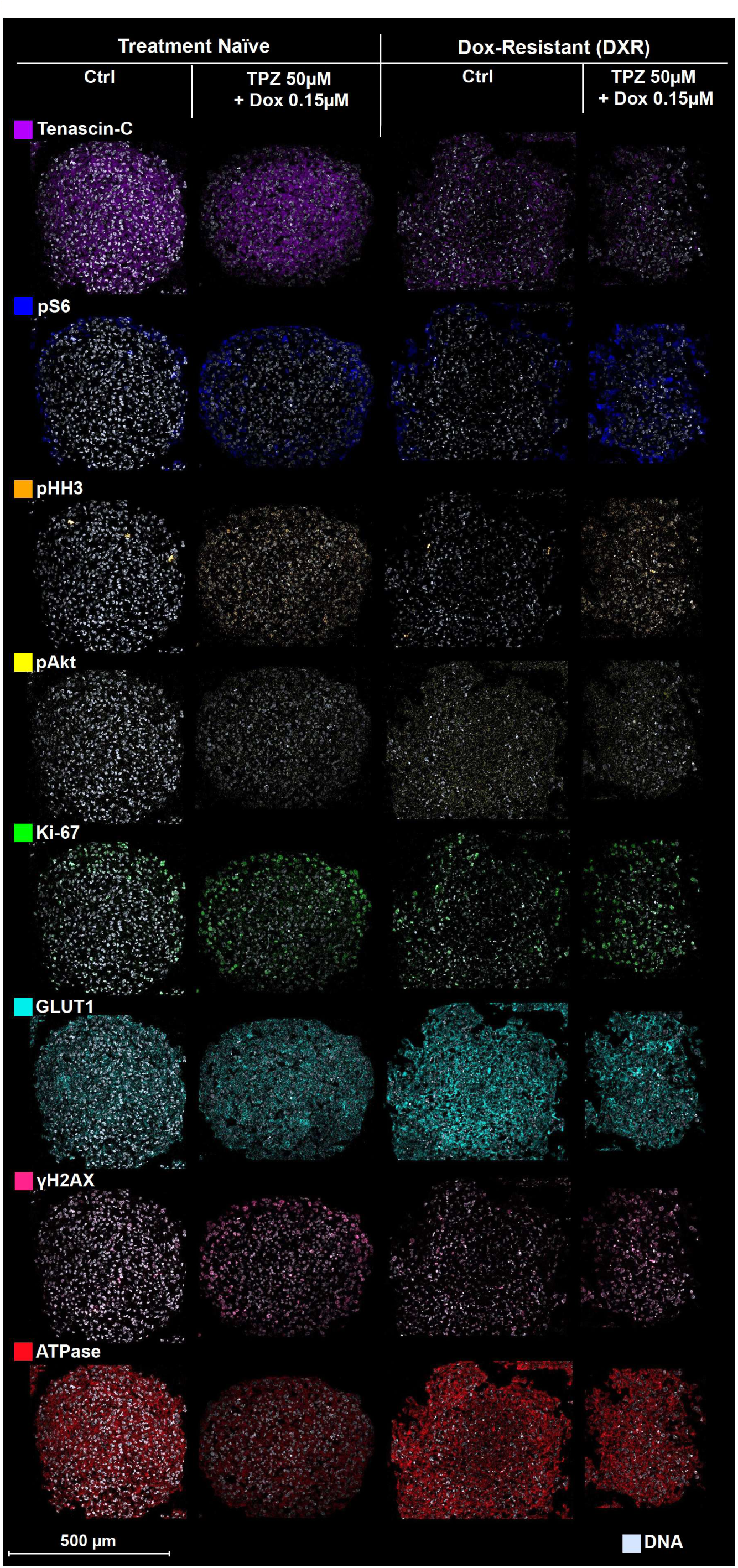
Spatial Protein analysis of MG-63 MCTS models across four treatment groups; treatment-naïve untreated (control), treatment-naïve acutely treated with TPZ-A-X 50μM/Dox 0.15μM, DXR untreated (control) and DXR acutely treated with TPZ-A-X 50μM/Dox 0.15μM. Protein localization analyzed at 1μm spatial resolution by IMC, shown by individual images of Tenascin-C (purple), pS6 (blue), pHH3 (orange), pAkt (yellow), Ki-67 (green), GLUT1 (cyan), γH2AX (pink), ATPase (red), and DNA (grey).

**Figure S3.**
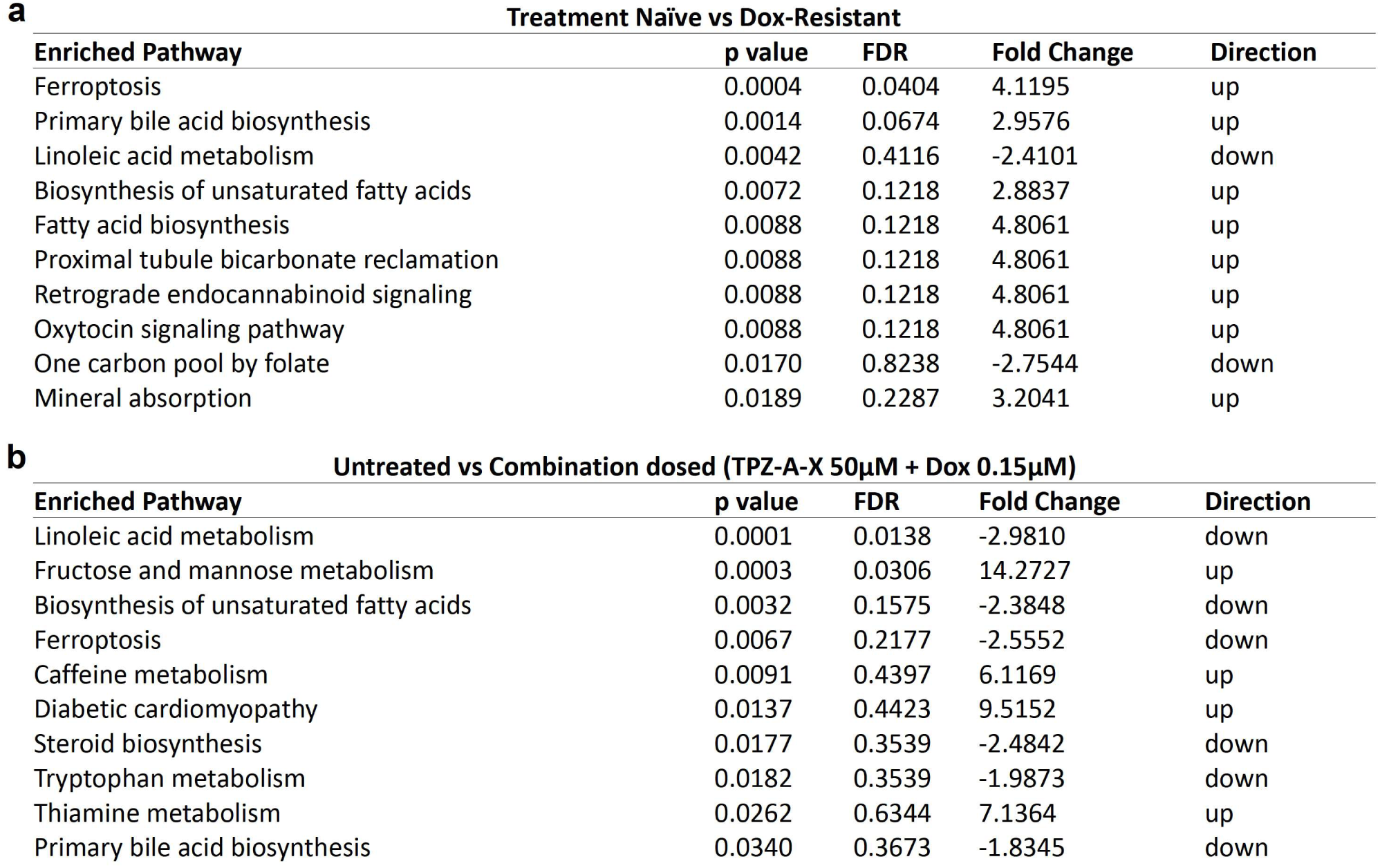
Top 10 pathway enrichment hits, sorted by raw p value, when comparing; (a) Treatment-naïve to DXR controls, as presented in the metabolic pathway enrichment volcano plot in Figure 3a, and (b) Acute combination treatment (TPZ-A-X 50μM/Dox 0.15μM) compared to untreated controls across both naïve and resistant treatment groups, presented in the metabolic pathway enrichment volcano plot in Figure 3b and c.

**Figure S4.**
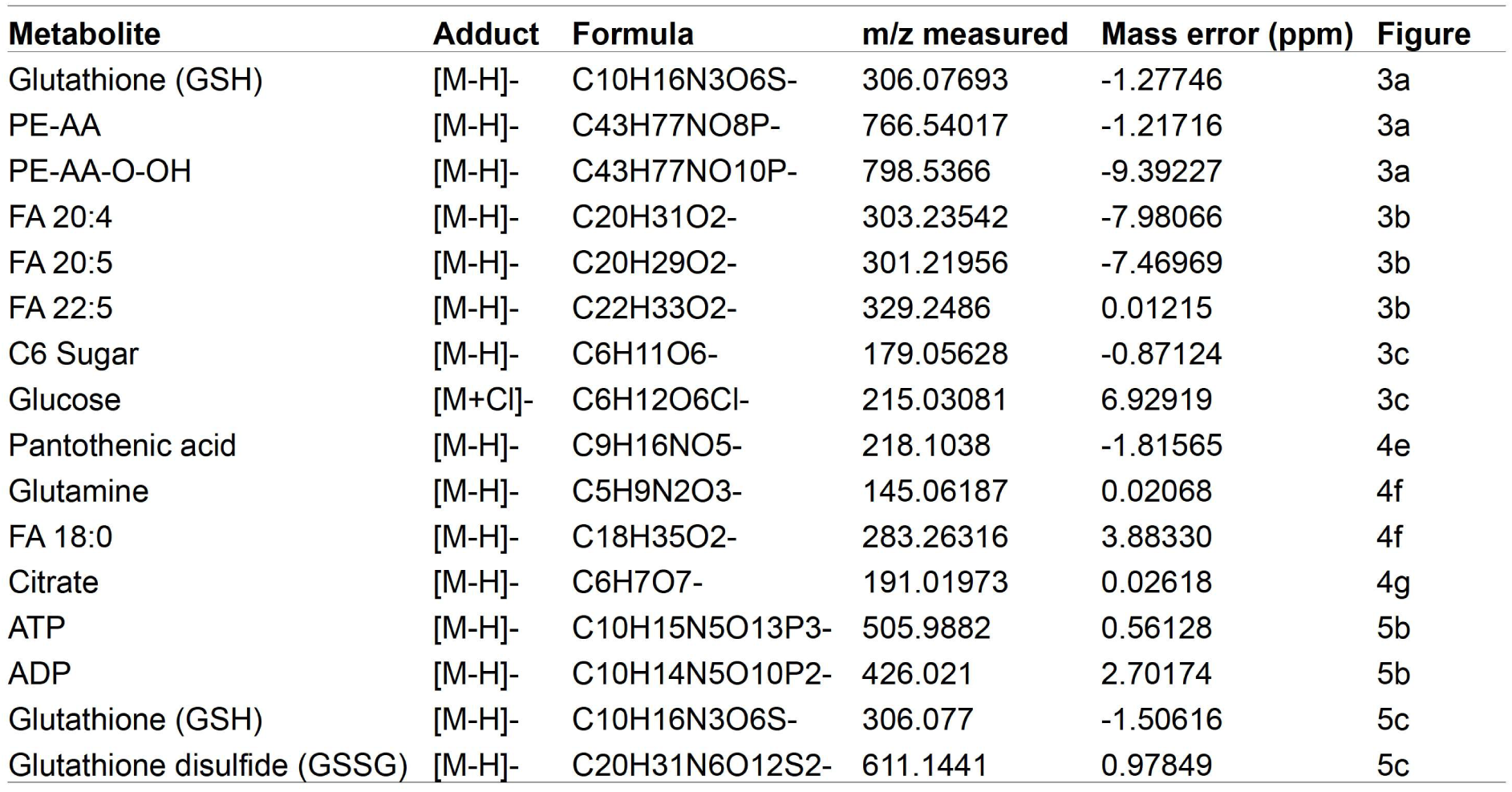
Annotated molecular ions and corresponding mass errors.

**Figure S5.**
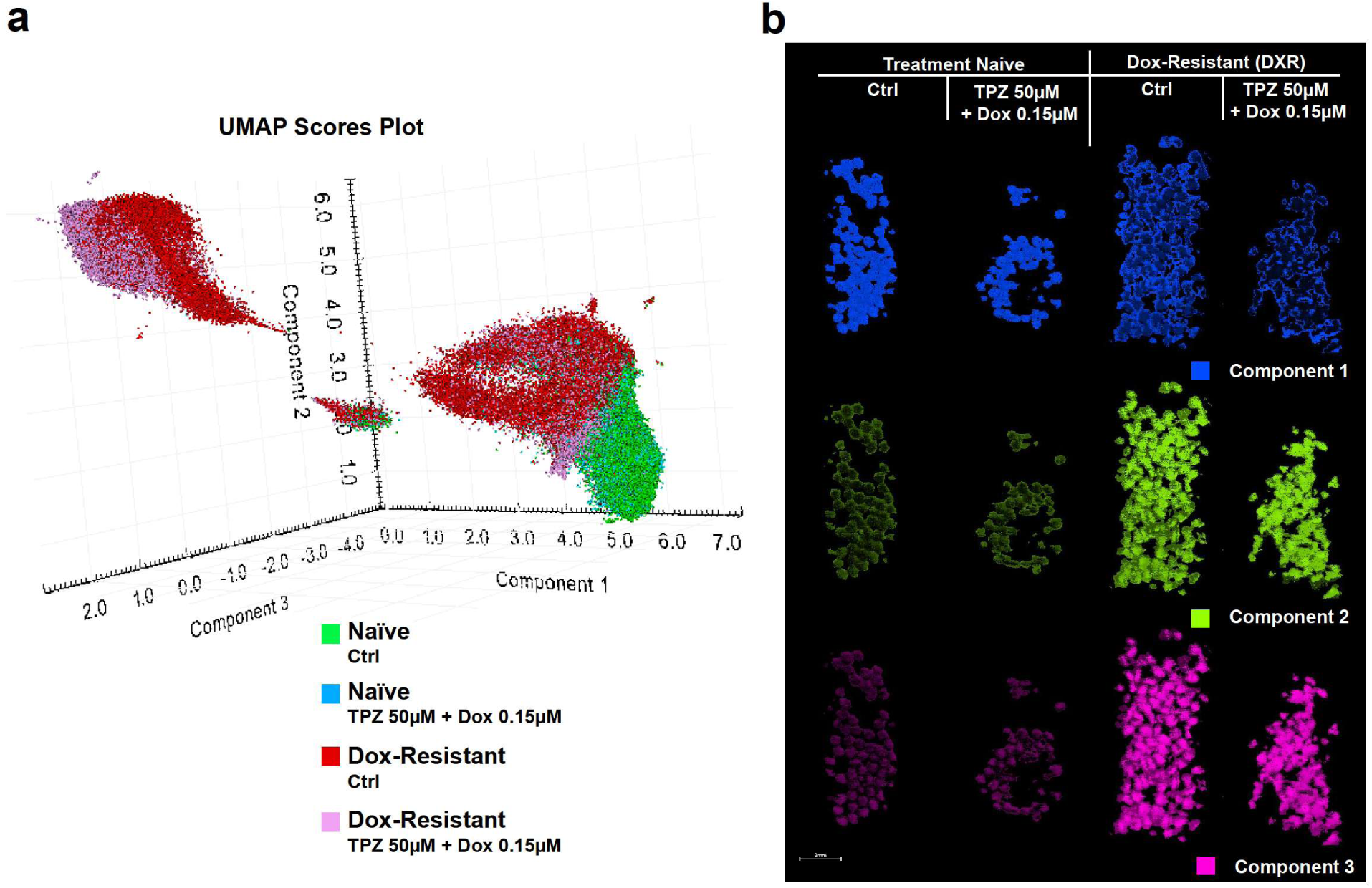
The Uniform Manifold Approximation and Projection (UMAP) components discussed in Figure 3 following spatial metabolomics by DESI-MSI, are here mapped onto the tissue regions. (a) The UMAP scores plot is shown, whereby scores plot points are color coded by treatment group: treatment-naïve untreated (green), treatment-naïve acutely treated with TPZ-A-X 50μM/Dox 0.15μM (blue), DXR untreated (red) and DXR acutely treated with TPZ-A-X 50μM/Dox 0.15μM (pink). (b) The identified components shown spatially mapped onto the analyzed tumor models. Component 1 is shown in blue, component 2 is shown in green, and component 3 is shown in pink.

**Figure S6.**
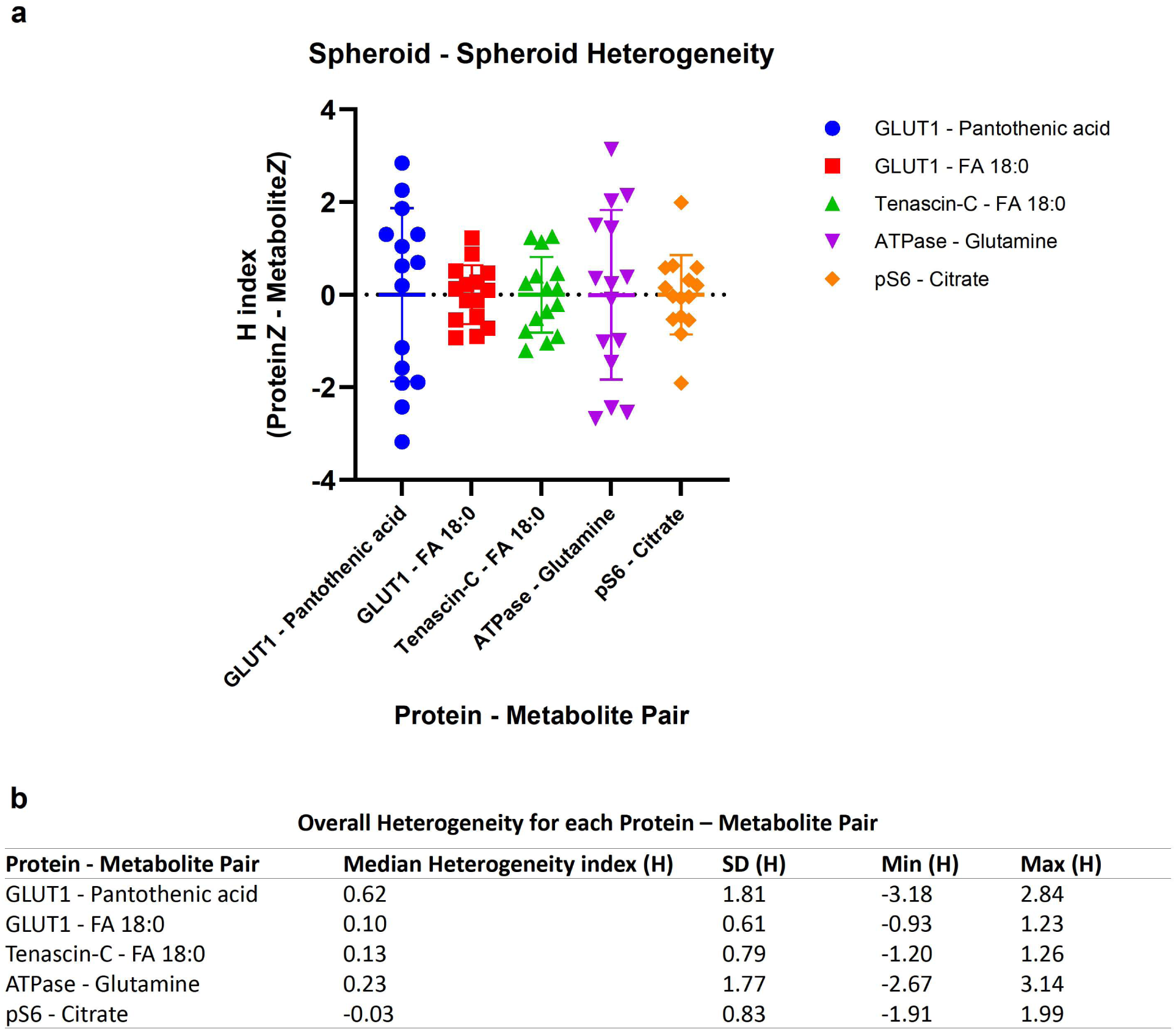
Individual tumor model heterogeneity of protein-metabolite correlations. (a) Distribution of spheroid to spheroid-level heterogeneity indices (H) for the five protein–metabolite pairs presented in Figure 4. Each point represents a single spheroid (n = 15 per pair). H index was calculated as the absolute difference between protein and metabolite z-scores within each spheroid, capturing the degree of spatial heterogeneity contributing to the observed correlations. Error bars represent standard deviation. (b) Summary statistics of the overall heterogeneity for each protein–metabolite pair, showing mean H ± SD alongside the minimum and maximum values across spheroids. Together, these data demonstrate that significant protein–metabolite correlations are driven by heterogeneous spheroid-level expression patterns.

**Figure S7.**
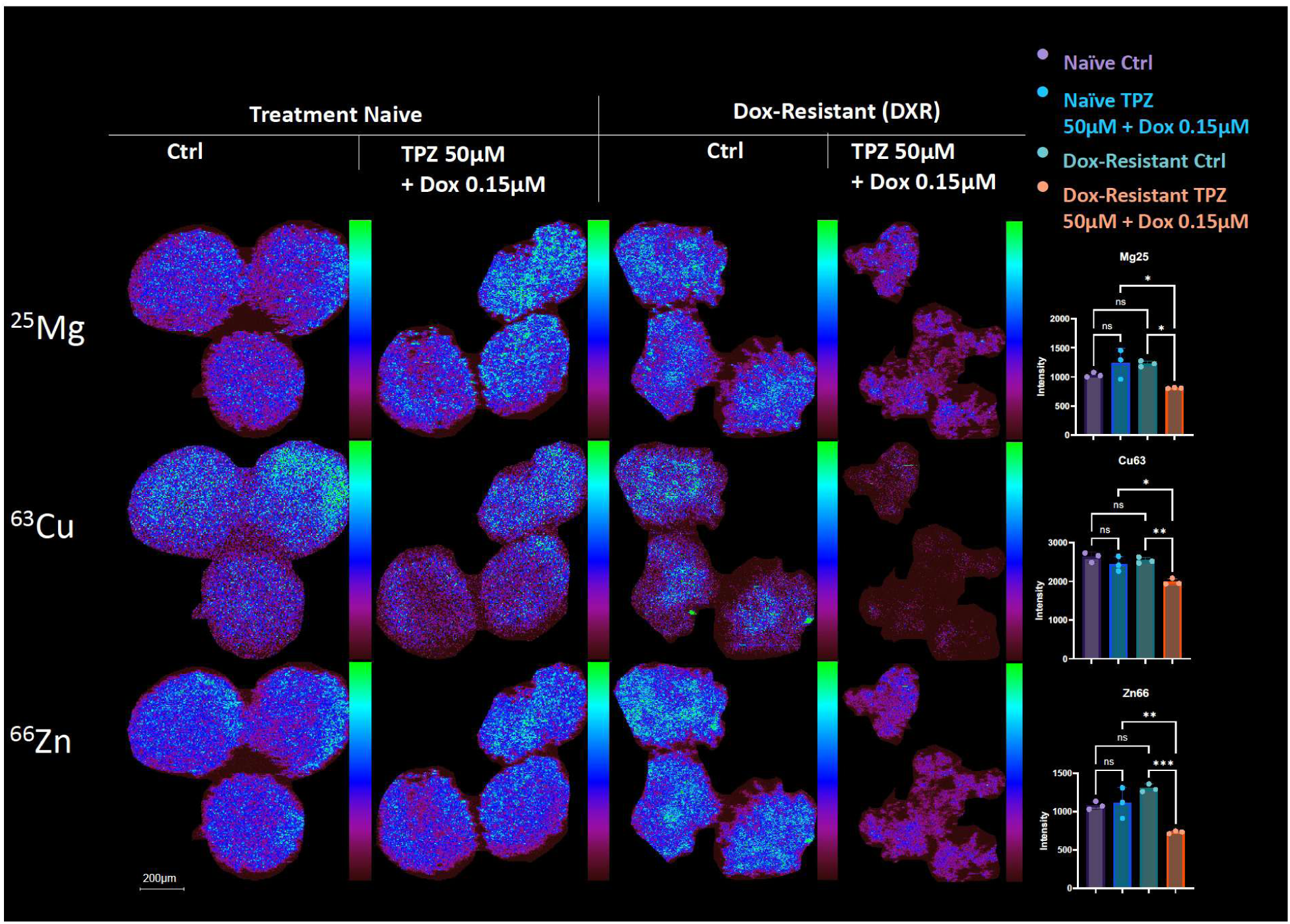
Endogenous metal isotope intensities shown to complement corresponding core:entire region ratios presented in Figure 5. Endogenous metal analysis by LA-ICP-MSI at 5μm spatial resolution. Ion images of magnesium (^25^Mg), copper (^63^Cu) and zinc (^66^Zn) are displayed across treatment groups, and their average intensities are shown. The data from the entire region of each spheroid were extracted to calculate average intensities Within each treatment group, 3 spheroids were analyzed (n = 3).

**Figure S8.**
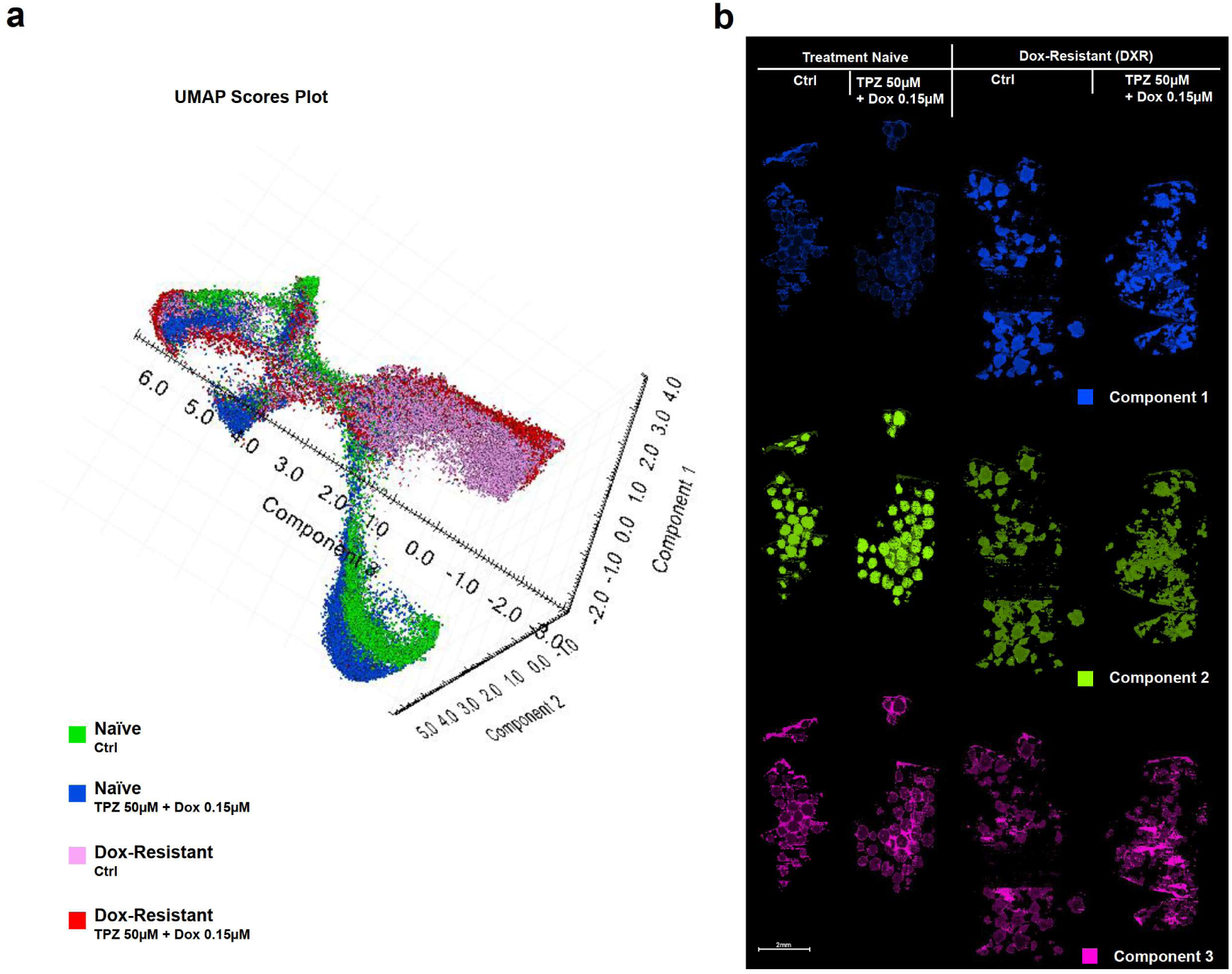
The Uniform Manifold Approximation and Projection (UMAP) components discussed in Figure 5 following spatial metabolomics by MALDI-MSI, are here mapped onto the tissue regions. (a) The UMAP scores plot is shown, whereby scores plot points are color coded by treatment group: treatment-naïve untreated (green), treatment-naïve acutely treated with TPZ-A-X 50μM/Dox 0.15μM (blue), DXR untreated (red) and DXR acutely treated with TPZ-A-X 50μM/Dox 0.15μM (pink). (b) The identified components shown spatially mapped onto the analyzed tumor models. Component 1 is shown in blue, component 2 is shown in green, and component 3 is shown in pink.

